# Gene expression profiles complement the analysis of genomic modifiers of the clinical onset of Huntington disease

**DOI:** 10.1101/699033

**Authors:** Galen E.B. Wright, Nicholas S. Caron, Bernard Ng, Lorenzo Casal, Xiaohong Xu, Jolene Ooi, Mahmoud A. Pouladi, Sara Mostafavi, Colin J.D. Ross, Michael R. Hayden

## Abstract

Huntington disease (HD) is a neurodegenerative disorder that is caused by a CAG repeat expansion in the *HTT* gene. In an attempt to identify genomic modifiers that contribute towards the age of onset of HD, we performed a transcriptome wide association study assessing heritable differences in genetically determined expression in diverse tissues, employing genome wide data from over 4,000 patients. This identified genes that showed evidence for colocalization and replication, with downstream functional validation being performed in isogenic HD stem cells and patient brains. Enrichment analyses detected associations with various biologically-relevant gene sets and striatal coexpression modules that are mediated by CAG length. Further, cortical coexpression modules that are relevant for HD onset were also associated with cognitive decline and HD-related traits in a longitudinal cohort. In summary, the combination of population-scale gene expression information with HD patient genomic data identified novel modifier genes for the disorder.

## INTRODUCTION

While genomic studies have made significant progress in identifying genetic variants associated with human disease, there has been less focus on the study of genomic modifiers of disease to date.^1, 2^ Such research is of great importance since it can be used to complement the development of genetic prediction models, provide novel therapeutic approaches and improve genetic counselling strategies. Huntington disease (HD) is an autosomal dominant neurodegenerative disorder that is caused by an expanded CAG tract in the *HTT* gene.^3^ Individuals with repeat lengths of 40 CAGs and greater display full penetrance and there is an inverse relationship between repeat length and clinical age of onset of the disorder.^4^ However, the length of this repeat only explains approximately 67% of the variability in age of onset observed between affected individuals and a large proportion of the residual clinical differences in age of onset observed between patients is likely to be heritable.^5, 6^ Identifying this residual component is important since HD is the most prevalent monogenic neurological condition in the developed world.^3^ Moreover, understanding this genetic component will help guide the selection of participants and objective measures for clinical trials evaluating experimental therapeutics.

Recent studies have started to make progress in identifying modifier genetic variants, both for HD,^7–11^ and other neurological disorders.^12–14^ For example, genetic variants in the interrupting sequence between the pathogenic CAG repeat and the polymorphic CCG repeat have been shown to influence age of onset of HD patients, and are particularly relevant for patients that carry reduced penetrance alleles.^10, 11^ Additionally, genome-wide association studies (GWAS) have identified *trans*-modifiers of clinical age of onset in HD,^7, 8, 10^ and have highlighted the important role of DNA repair genes in modulating age of onset, potentially through altering the somatic instability of the CAG repeat.

Interestingly, a subset of these HD modifiers has been shown to modulate the age of onset of spinocerebellar ataxias, which are also caused by pathogenic CAG repeats,^15^ indicating the potential transferability of these genomic findings in HD to related disorders. While these studies have made significant advances in our understanding of the genetic modulation of clinical onset in HD, they only explain a limited proportion of the residual variance observed between HD patients. Further, since the signals derived from GWAS are typically in non-coding regions, prioritized regions still need to be refined through the incorporation of functional genomic information into analyses.^16^ This is pertinent, since it is well documented that changes in gene expression are a hallmark of HD disease pathology,^17, 18^ and *HTT* is expressed ubiquitously across tissues.^19^

Examination of heritable gene expression profiles has recently been made possible through the development of transcriptome-wide association study (TWAS)-based approaches. These enhance GWAS-derived information through leveraging population-scale expression quantitative trait locus (eQTL) data derived from diverse tissues.^20–22^ These analyses are used to assess the relevance of gene expression with regards to the GWAS trait, returning gene-level association results. Due to a reduced multiple testing burden compared to GWAS, these transcriptome-inferred analyses have more power to prioritize additional candidate regions, which can in turn be used to inform future *in vitro* and *in vivo* genetic modifier validation studies. Further, gene-level association results can also be used in enrichment analyses to determine whether particular pathways or gene sets are relevant to the trait of interest.

We therefore assessed whether heritable variation in gene expression was associated with the clinical onset of HD by analyzing GWAS data for this phenotype from over 4,000 patients in combination with transcriptomic information from 48 tissues (summarized in Fig. 1). The expression of prioritized genes was then further evaluated in an allelic series of isogenic HD human pluripotent stem cells (hPSCs), as well as through an orthogonal analysis of proteins in pathologically-relevant brain regions in HD patient samples. Building on these findings, we showed an enrichment for associations in disease-relevant gene sets, as well as in genes belonging to coexpression modules that are either huntingtin-CAG dependent or associated with disease-relevant traits in the human brain. Finally, we performed HD TWAS signature matching to inform future studies of drug repurposing.

**Fig. 1.**
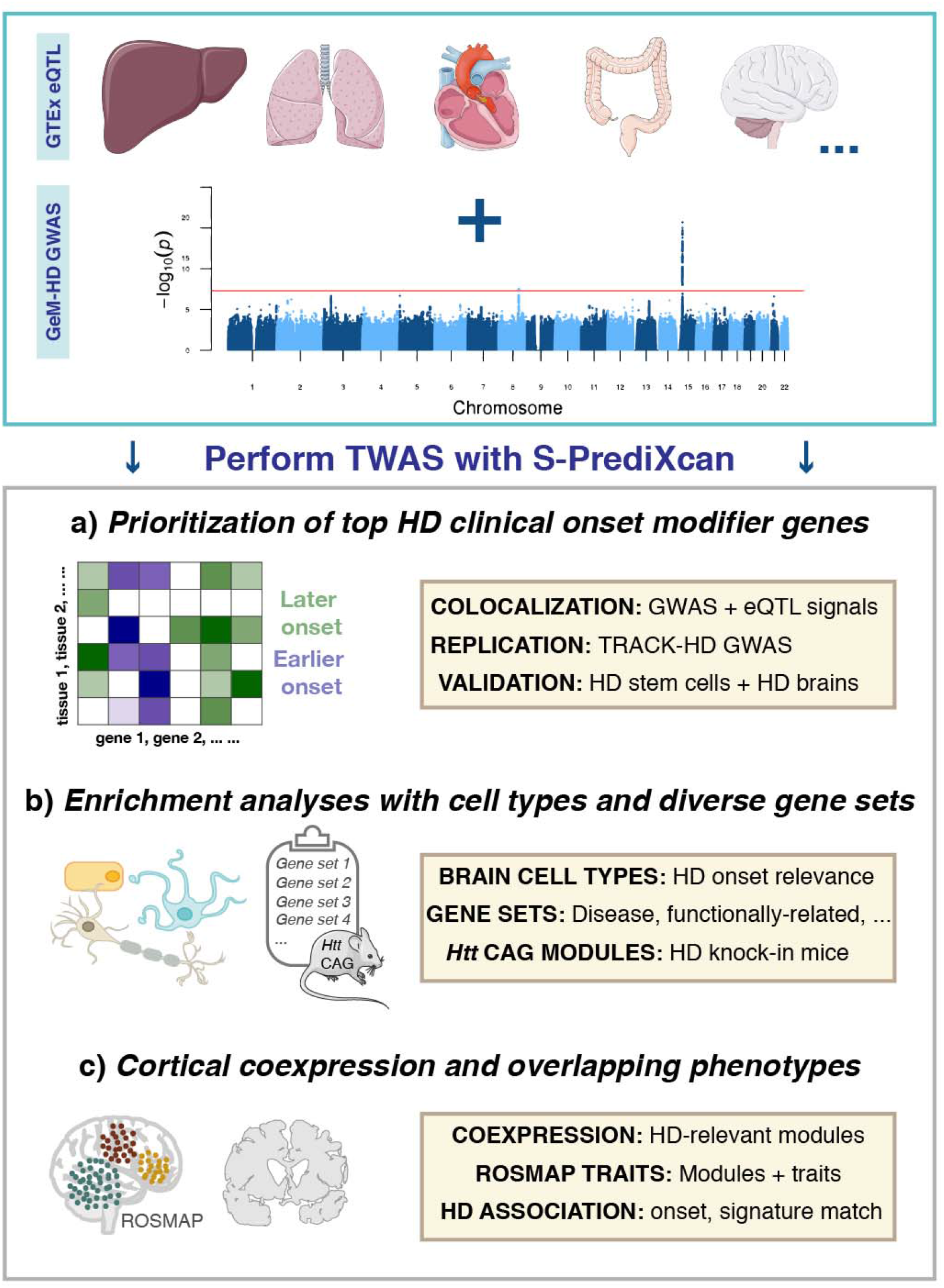
Schematic representation methodology employed to identify transcriptomic modifiers of Huntington disease, as well as to gain additional insights into the biological mechanisms that underlie this trait. Transcriptome wide association analyses facilitated **(a)** the identification of top transcriptomic HD onset modifier genes, **(b)** relevant biologically diverse gene sets that are enriched for this trait, and **(c)** human cortical coexpression modules that are relevant for HD onset and related phenotypes in an aging cohort, along with signature matching of HD signals for drug repurposing.

## RESULTS

### Transcriptomic analyses prioritize novel genomic modifiers for the clinical onset of HD

Since mutant *HTT* is expressed throughout the body and induces pathology in numerous tissues,^3^ we chose to perform an agnostic approach to tissue selection, using all information available from the Genotype-Tissue Expression (GTEx) Project. This approach is also recommended by the S-PrediXcan developers to maximize discovery power.^20^ Combining these eQTL data with the HD clinical onset GWAS, TWAS in the discovery cohort identified 15 candidate genes located in eight genomic regions (Table 1, Fig. 2, Supplementary Fig. 1 and Supplementary Table 1) where the genetic component of expression was predicted to significantly associated with HD clinical onset.

**Table 1.**
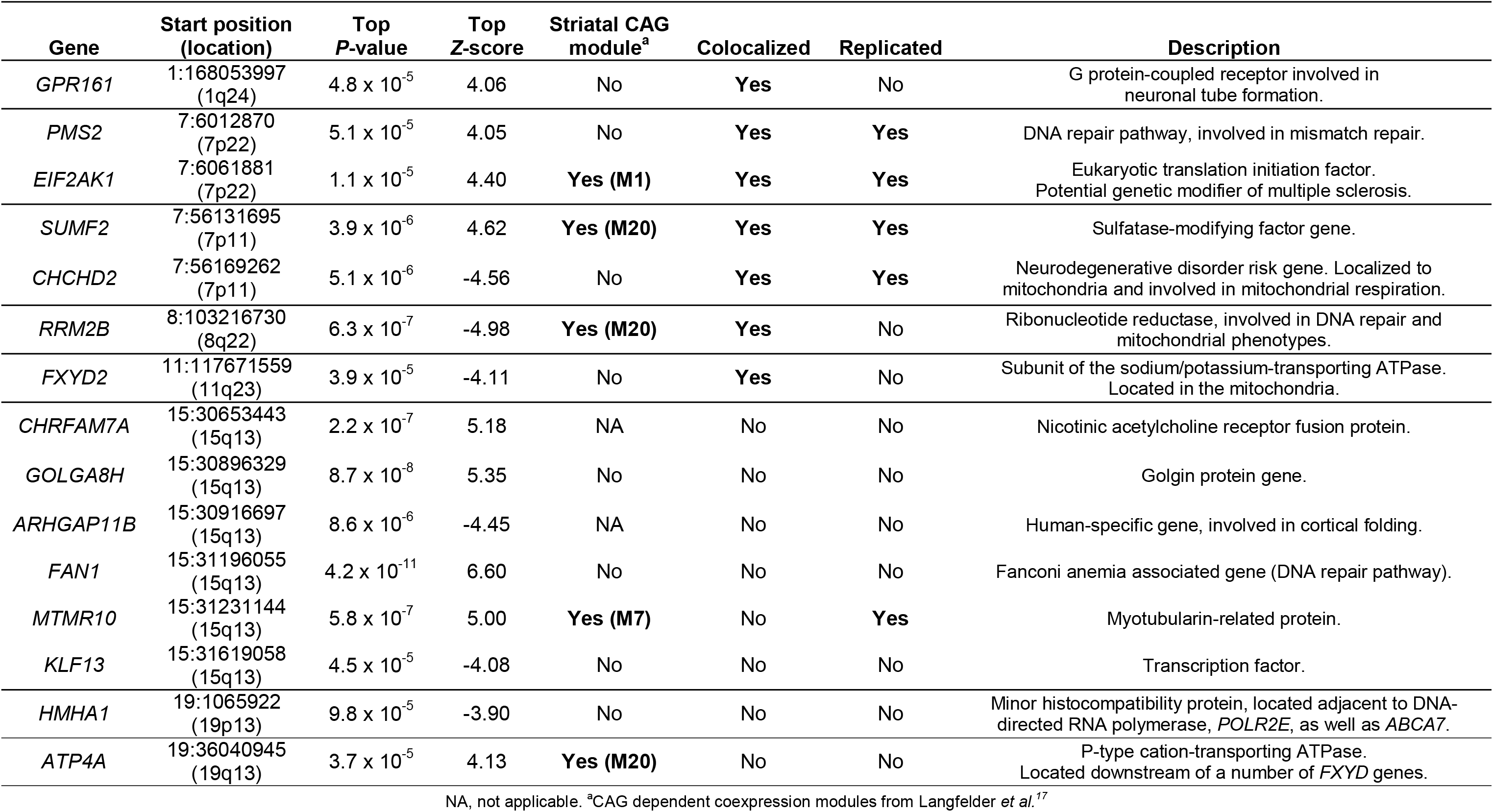
Summary of the 15 prioritized transcriptomic modifier genes where imputed expression is associated with the age of clinical onset of Huntington disease in the GeM-HD cohort. Genes are annotated for chromosomal location, whether the expression and GWAS based signals colocalize, whether there was evidence for replication in the TRACK-HD cohort and membership to striatal coexpression modules where expression is influenced by *Htt* CAG repeat lengths.

**Fig. 2.**
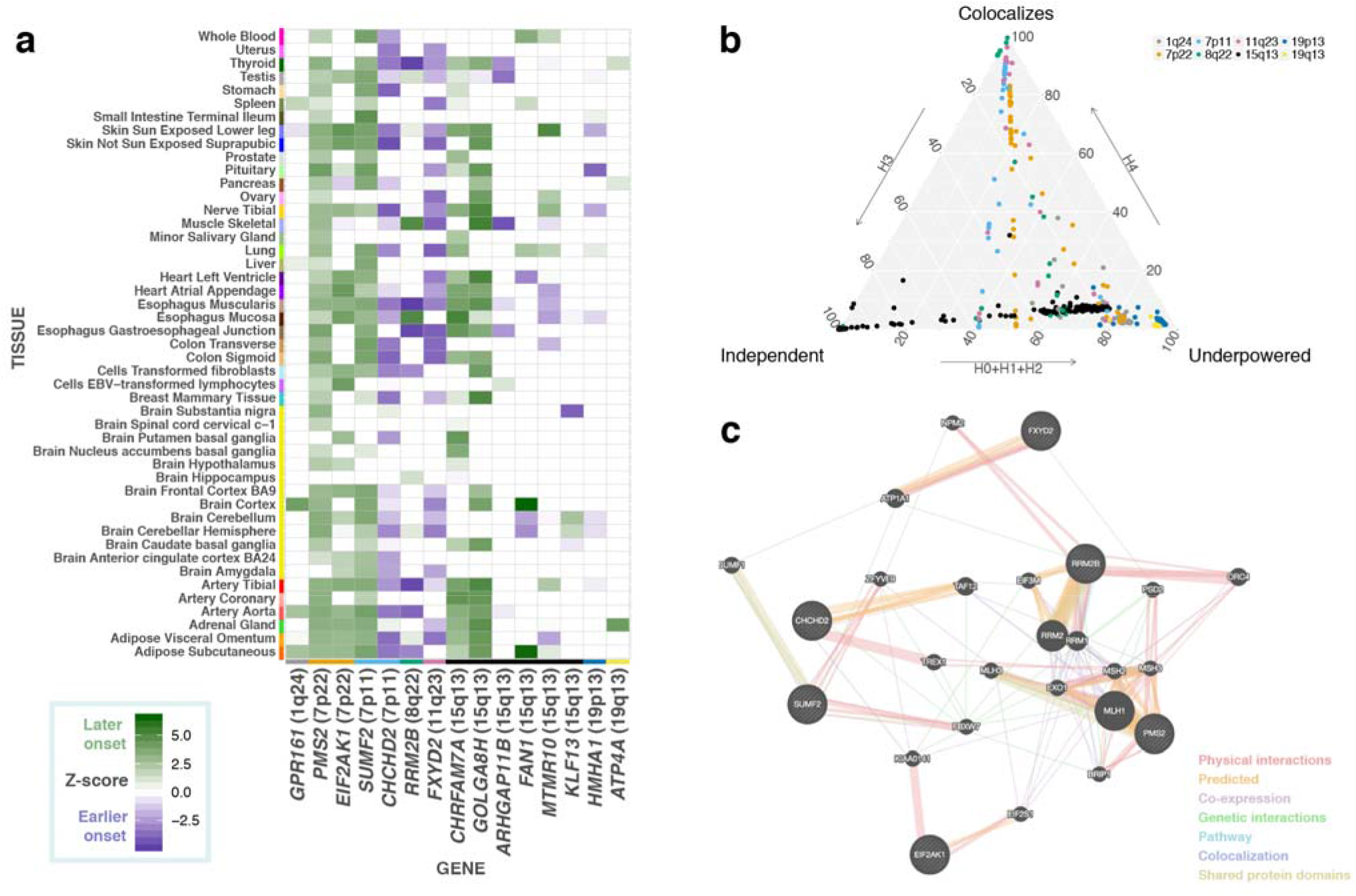
Transcriptome wide association analyses of Huntington disease clinical onset prioritizes novel candidate modifier genes. **(a)** Gene level *Z*-scores of top modifier genes across tissues in the GeM-HD discovery cohort. Positive *Z*-scores indicate that increased gene expression is associated with a later age of clinical onset in patients, while negative *Z*-scores are associated with earlier onset. **(b)** Ternary plot of COLOC posterior probabilities colored by chromosomal location displaying TWAS regions that displayed evidence for colocalization (i.e. COLOC *P4*>0.5) versus those regions that were either underpowered or where eQTL and GWAS signals represent independent associations. **(c)** These colocalization analyses aided in refining Huntington disease modifier gene list as indicated by the tight connections observed in the gene-gene functional interaction network that was enriched for mismatch repair, DNA binding and ATPase activity. *GPR161* is the only colocalized gene that does not form part of this network.

The most significant TWAS association was for *FAN1* in the cortex (*Z*-score 6.60, *P*=4.25 × 10^−11^), a signal which appears to be driven by eQTLs linked to the one of the chromosome 15 signals detected in the GeM-HD GWAS (top GWAS variant: rs2140734, delays onset by 1.4 years, GTEx eQTL cortex: normalized effect size 0.38, *P*=3.2 × 10^−7^). However, examination of the top GWAS variant in a larger cortical dataset,^23^ revealed that this variant is most significantly associated with *MTMR10* expression (Spearman’s rank correlation=0.24, *P*=8.51 × 10^−8^). Four novel prioritized genes at two loci (*PMS2 P*=5.1 × 10^−5^; *EIF2AK1 P*=1.1 × 10^−5^; *SUMF2 P*=3.9 × 10^−6^ and *CHCHD2 P*=5.1 × 10^−6^), showed evidence for colocalization between the GWAS and expression association signals (Supplementary Table 2), as well as independent replication in the TRACK-HD analyses. Of note, protein truncating mutations in the mismatch repair gene, *PMS2*, have been shown to cause cancers such as Lynch syndrome.^24^ Increased expression of *PMS2* (*Z*-score=4.05) was associated with later age of HD onset and could potentially play a role in mediating the somatic instability of the CAG repeat. The *EIF2AK1* and *CHCHD2* genes are relevant for neurological phenotypes, since genetic variants have been respectively associated with modulating multiple sclerosis^12^ progression, as well as risk for Parkinson disease and Lewy body disorders respectively.^25^

The association of multiple genes at the same locus, supported by colocalization analyses, could be the result of co-regulation. Colocalization analyses employ a Bayesian statistical test to assess whether two genomic association signals share a common causal variant. They therefore serve as a post-filtering step to prioritize signals where there is a high probability that the GWAS and eQTLs association results arise from a single causal variant, as opposed to S-PrediXcan associations that may have resulted simply from linkage disequilibrium (Fig. 2, Supplementary Table 2). The value of these colocalization analyses is indicated by the clear clustering of colocalized modifier genes into biological networks, including an enrichment for DNA repair genes with ATPase activity (Fig. 2). It should, however, be noted that non-colocalization could also have been observed for some loci due to limited power, or multiple causal variants at a chromosomal region. None of the genes from the top GWAS signal on chromosome 15 were colocalized, which may reflect the COLOC model assumption of one causal variant, which reduces the accuracy of these outputs when multiple causal variants are observed.^26^

Examination of the remaining TWAS signals uncovered additional noteworthy genes. These included *RRM2B*, a gene involved in DNA replication and repair; and *GPR161*, a cilium-related G protein-coupled receptor that has been implicated in neural tube development via the sonic hedgehog pathway^27^, both of which displayed evidence of colocalization. Further, novel associations with ATPase-related genes, *ATP4A* and *FXYD2*, were also detected, which could be of relevance to other findings since many DNA repair proteins possess conserved ATPase domains.^28^ The Rho GTPase activating protein gene, *HMHA1*, was prioritized on chromosome 19p13, however, this gene is located directly downstream of a notable Alzheimer’s disease gene,^29^ *ABCA7*. Although *ABCA7* was not prioritized after accounting for multiple testing across tissues, over 60% of *ABCA7*-tissue combinations displayed nominally significant *P*-values, indicating that variants related to this gene may have caused this signal. Finally, the mismatch repair gene, *MSH3* gene, which was prioritized in the TRACK-HD GWAS^7^ used in our replication analyses, also displayed a number of highly-ranked associations in the discovery TWAS (top *Z*-score=-3.75, *P*=1.77 × 10^−4^).

### Further assessment of prioritized transcriptomic HD modifier genes in model systems and HD patient brains

We assessed alterations of top candidate transcriptomic modifiers at the gene expression level in an allelic series of isogenic HD hPSCs, and at the protein level in cortical and striatal HD patient and age-matched control brains. Using isogenic HD hPSCs as an *in vitro* model of HD, it was found that DNA repair genes (i.e. *FAN1, PMS2* and *MSH3)* displayed increased expression levels at longer CAG repeat lengths in pluripotent stem cells (Fig. 3). Cells in this early developmental stage display increased activity of DNA repair systems,^30^ and the higher expression levels at larger CAG lengths could be the result of preventative mechanisms relating to the repeat expansion. Conversely, protein levels for these genes were decreased in HD patient brains compared to controls (Fig. 3 and Supplementary Table 3; *FAN1* cortex *P*=0.0004; *PMS2* cortex *P*=0.017; *MSH3* caudate nucleus *P*=0.04). This may therefore reflect deficiencies in DNA repair in the final stages of the disease. Alternatively, negative correlations between RNA and protein levels have been previously observed for a subset of genes in knock-in *mHtt* mice, which may reflect the consequences of impaired post-transcriptional regulation as a result of mutant huntingtin.^17^

**Fig. 3.**
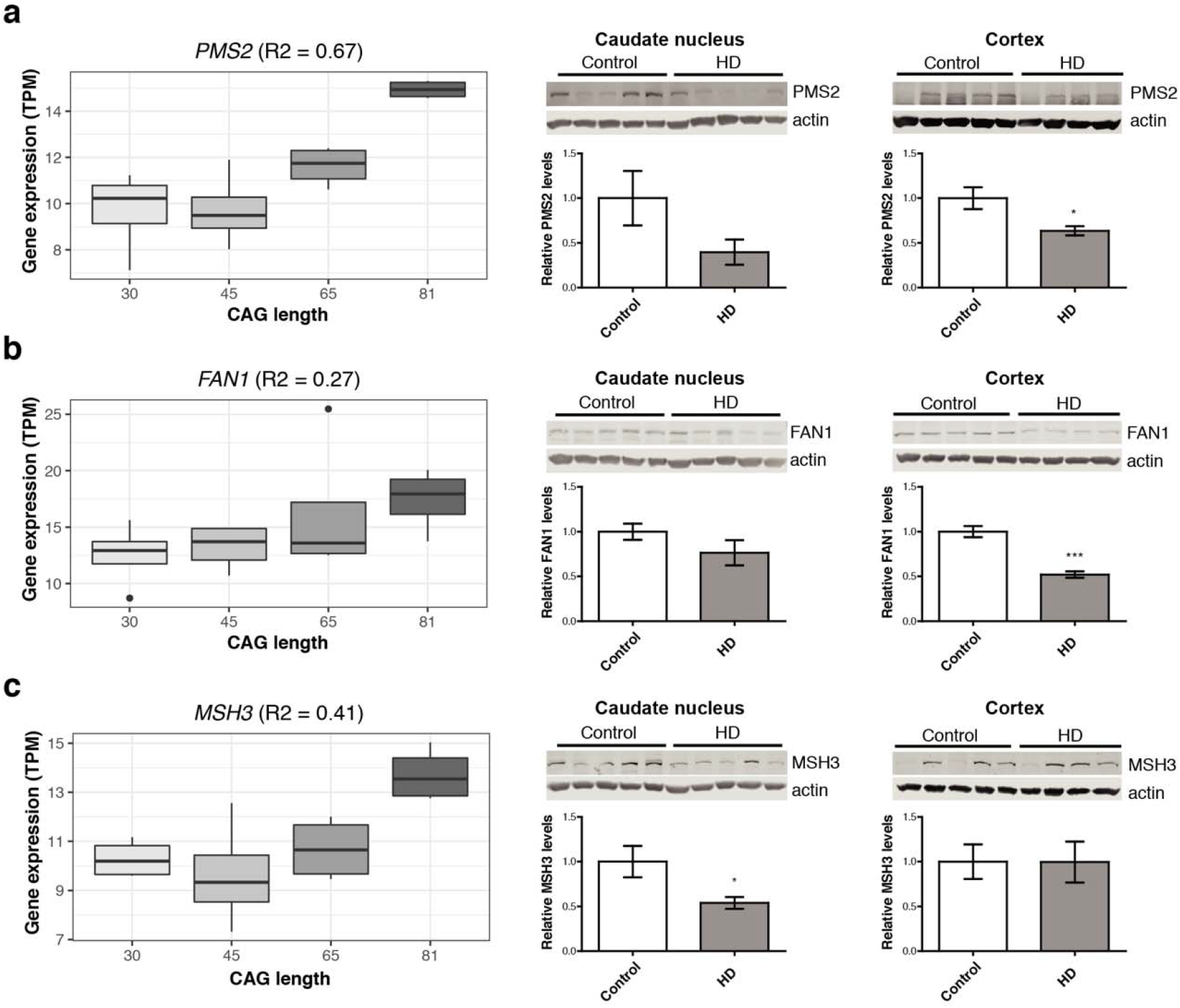
DNA repair genes are dysregulated in HD stem cell lines, as well as patient brains, but show differing directions of effect at the transcript and protein level. Gene expression of DNA repair genes, **(a)** *PMS2*, **(b)** *FAN1* and **(c)** *MSH3*, was elevated at longer CAG lengths in human pluripotent stems cells (hPSCs, left), but relative protein levels are decreased in HD patient brains compared to matched controls (middle and right). Asterisks indicate *P*<0.05.

Additional supporting evidence was also obtained *in vitro* for the *GPR161* gene, since expression levels were decreased at high CAG lengths in allelic neural progenitor cells (*R*^2^=0.43, *P*=0.006). Further, GPR161 protein levels were also lower in the caudate nucleus of HD patients compared to age-matched controls (*P*=0.005, Fig. 4), with a trend towards significance in the same direction in the cortex in these patients (Supplementary Table 3). The most significant TWAS association for *GPR161* was in the cortex, indicating that increased gene expression is associated with a later age of onset in HD patients. This influence on HD clinical onset is in line with previous experiments that have observed increased genomic instability of medulloblastomas in conditional knockout *Gpr161* mice.^27^ Further, the recent GeM-HD GWAS preprint^10^ has implicated genetic variation in another G protein-coupled receptor gene, *GPR151*, in modifying clinical onset in HD. Network-based analyses of *GPR161* and *GPR151* suggest the importance of serotonin signaling through shared protein domains of the receptors (Supplementary Fig. 2). These analyses also reveal potential interactions between *GPR161* and the melanocortin 2 receptor gene, *MC2R*.

**Fig. 4.**
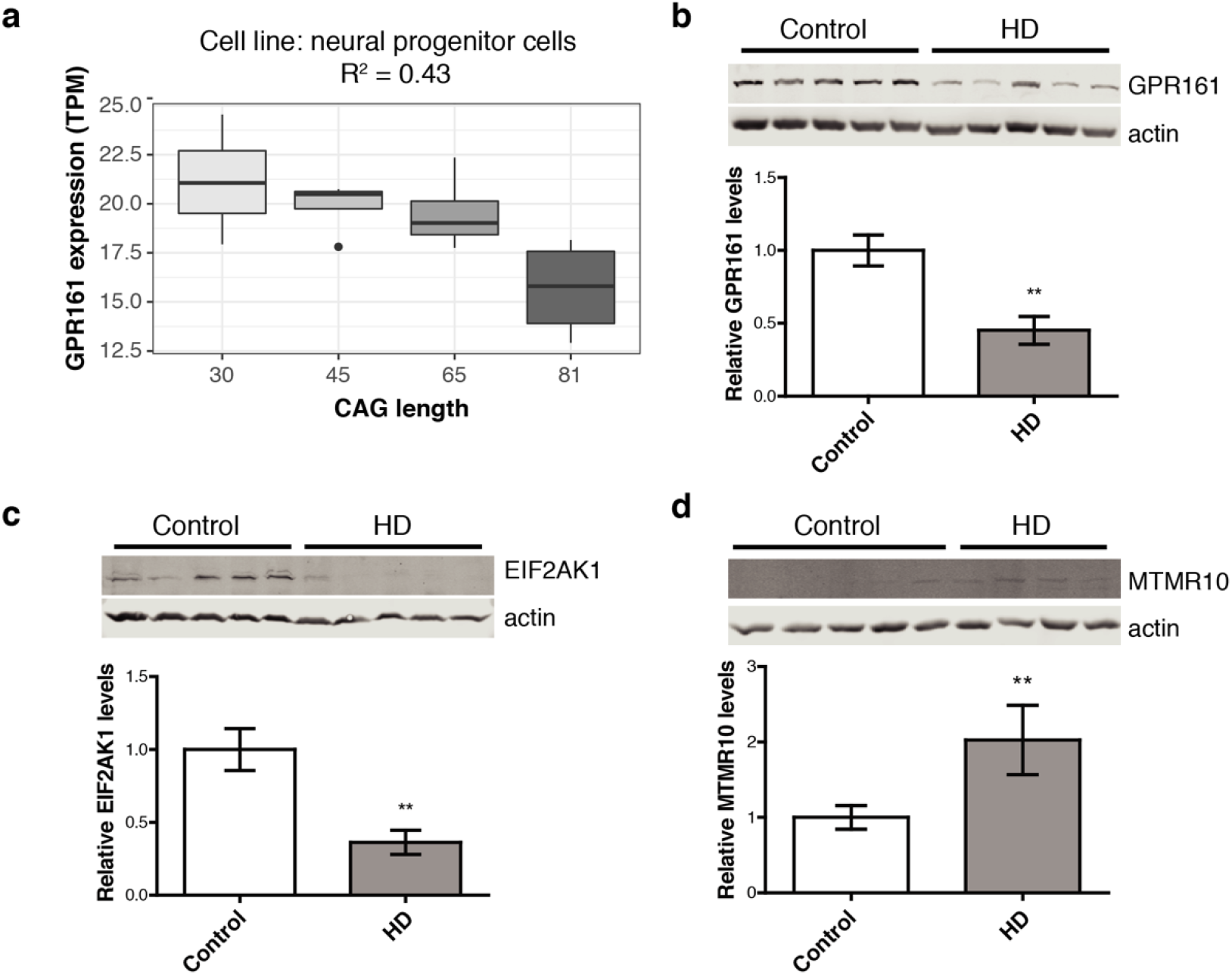
Orthogonal validation of key transcriptomic modifier genes confirms the dysregulation of candidate HD modifiers in humans. The ciliary G-protein-coupled receptor, *GPR161*, was **(a)** downregulated at increasing CAG lengths in neural progenitor cells and **(b)** decreased relative protein levels in the caudate nucleus in HD patients compared to controls. Other candidate HD modifier proteins that were also dysregulated in this striatal region in patients include **(c)** EIF2AK1 and **(d)** MTMR10. Asterisks indicate *P*<0.05.

### HD transcriptomic modifier associations are enriched in relevant cell types and gene sets

Next, we searched for trait-relevant cell types in the brain for HD clinical onset using cortical single cell sequencing data. These analyses detected a significant relationship with brain microvascular endothelial cells (BMECs) and disease onset (*P*<0.05, Fig. 5). BMECs constitute the major structural components of the blood-brain barrier, and recent studies of iPSC-derived versions of this cell type have shown that HD BMECs display both angiogenic and blood-brain barrier impairments in HD patients compared to control lines.^31^ An analysis of the prioritized transcriptomic HD modifier genes in this published gene expression dataset revealed that the two colocalized and replicated chromosome 7p11 genes were dysregulated in BMECs derived from HD patients (i.e. *CHCHD2 P*=4.4 × 10^−9^, fold change −1.9; and *SUMF2 P*=0.0016, fold change 1.1). On the protein level, MSH3, RRM2B and EIF2AK1 were dysregulated in our HD BMEC cell lines (Supplementary Fig. 3, Supplementary Table 4), indicating the potential importance of these proteins in this cell type in HD.

In an attempt to identify groups of genes that are important for modifying HD clinical onset, we evaluated 17 diverse gene sets for enrichment with regards to TWAS associations (Fig. 5, Supplementary Table 5). There was a significant underrepresentation of GWAS catalogue genes, which reflects the fact that the majority of GWAS conducted to date focused on complex diseases, and that clinical modifiers of a Mendelian disease represents a unique phenotype. Disease-related genes with annotations in ClinVar^32^ were significantly enriched for HD-related associations, particularly those causing autosomal dominant disorders and ones that result from haploinsufficiency. Autosomal recessive genes, however, were not significantly enriched for HD clinical onset associations. These findings indicate that more subtle changes in expression of genes that are intolerant to loss of function variants may play an important role in modulating clinical age of onset in HD. Associations for DNA repair genes were also significantly enriched, adding confirmatory evidence for somatic instability as a modifier for HD clinical onset.^7, 33^ Of potential translational relevance, drug target genes^34^ were also prioritized in the analyses. These gene set analyses therefore provide an avenue to identify the biological mechanisms the underlie the differences in age of onset observed between HD patients.

**Fig. 5.**
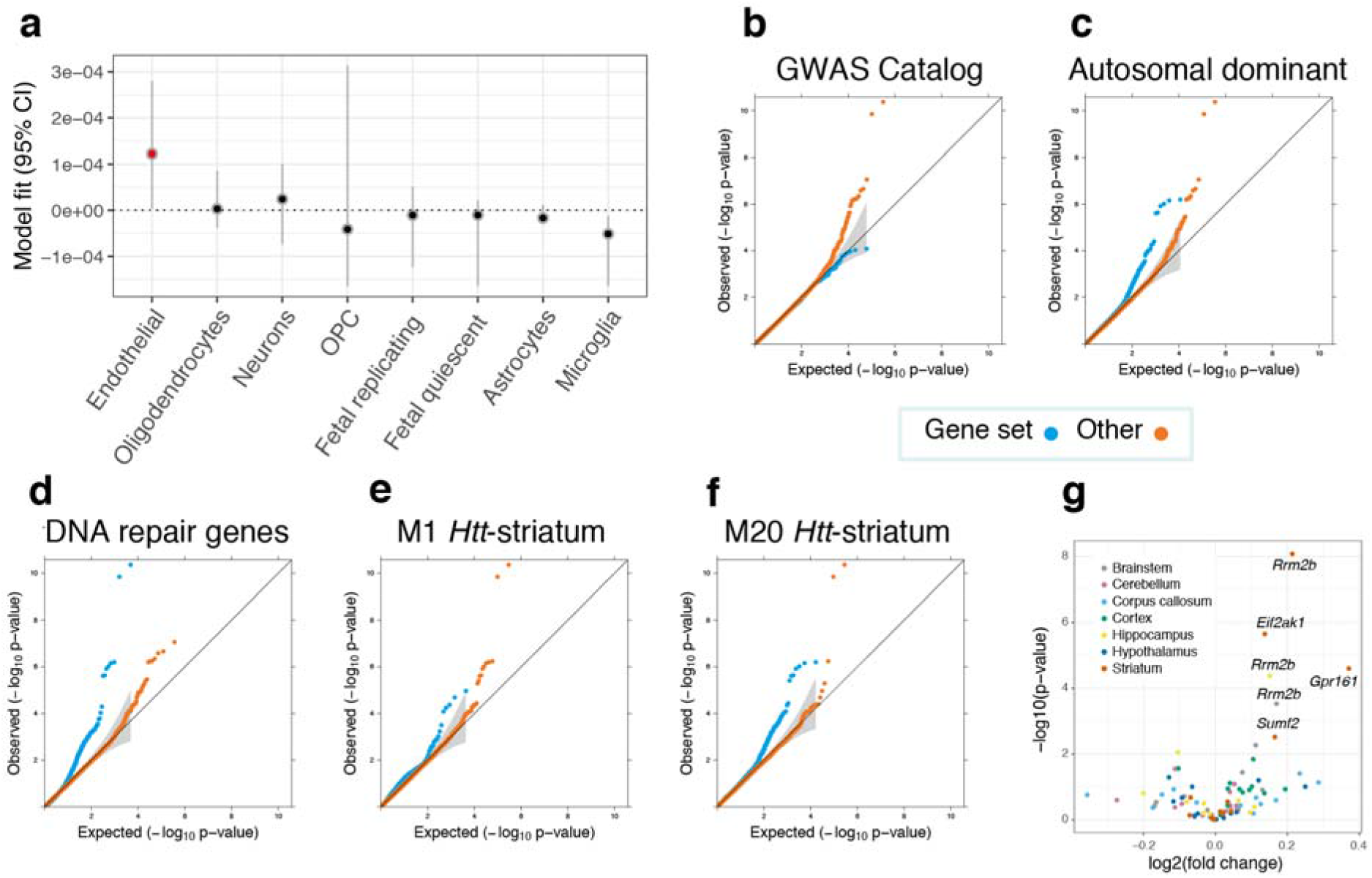
Enrichment analyses revealed brain cell types, striatal coexpression modules and gene sets that are most relevant for modifying the clinical onset of Huntington disease. **(a)** Inferring relevant brain cell types from the GWAS data supplemented with cortical single cell sequencing information suggests that brain microvascular endothelial cells (annotated in red) could be relevant for the trait (*P*<0.05). Cell types are ranked according to parameter estimates. Representative Q-Q plots for significantly enriched or depleted S-PrediXcan modifier associations for genes belonging to: various gene sets (**b** to **d**), as well as coexpression networks that are related to murine *Htt*-CAG size over time (**e** and **f**). M20_*Htt*-striatum_ genes are involved in p53 signaling and *Brca1* in DNA damage response, while M1_*Htt*-striatum_ genes are implicated in protein ubiquitination, tRNA charging and Huntington disease signaling. **(g)** Volcano plot for prioritized transcriptomic modifier genes in in Q175 mice versus wild type at 6 months of age for six brain-related tissues. Significantly dysregulated genes after correcting for multiple testing corrected are annotated, with all genes differentially expressed in the brain were upregulated in mice with expanded *Htt* CAGs.

### Gene coexpression modules that display a positive correlation to huntingtin CAG-length are enriched for HD genomic modifier associations

In order to determine whether groups of genes whose expression is mediated by mutant huntingtin were enriched in the HD modifier association results, we analyzed transcriptomic coexpression networks from a large time course experiment from HD knock-in mice with increasing *Htt* CAG lengths.^17^ An analysis of the 13 striatal and five repeat-length dependent cortical modules from Langfelder *et al.*^17^ revealed that two striatal modules, M1_*Htt*-striatum_ and M20_*Htt*-striatum_, were significantly enriched for S-PrediXcan associations (Supplementary Table 6). Both of these are positively correlated with *Htt*-CAG length. Interestingly, M20_*Htt*-striatum_ displayed the most significant association of these upregulated modules in the original study.^17^

### Analyses of gene expression profiles in the aging human brain identifies HD-relevant coexpression modules and overlapping phenotypes

Next, we assessed gene coexpression modules detected in the human cortex from a longitudinal aging cohort^35^ for enrichment in the HD-clinical onset TWAS results. This was performed in order to identify groups of co-expressed cortical genes that could be particularly important for modulating HD onset. Fifteen modules showed significant differences in mean *Z^2^*-scores between gene sets belonging to the cortical modules, with seven modules displaying a significant enrichment for HD-related phenotype associations compared to other genes (Fig. 6, Supplementary Table 7 and 8). Notably, the top m109_human-cortex_ module, that has been associated with Alzheimer’s disease and cognitive decline,^35^ was statistically underrepresented in HD-related associations (*P*=1.1 × 10^−11^). Conversely, the mitochondrial module, m131_human-cortex_, which was also implicated in cognition in that previous study, had significantly enriched HD-related scores when compared to other genes (*P*=4.1 × 10^−6^).

The ROSMAP study longitudinally collected various phenotypic measures for participants. This allowed us to assess which of these traits the seven HD onset-related coexpression modules are associated with in aging individuals, to inform what other biological processes these groups of genes are important for. In these analyses, increased expression of m122_human-cortex_ and m131_human-cortex_ was associated with improved performance for cognitive-related traits, while the opposite effect was observed for m108_human-cortex_ and m114_human-cortex_ expression (Fig. 6). Beneficial cortical modules were enriched for mitochondrial (m106, m122 and m131) and neuronal/synaptic (m16) gene ontology terms, while those deleterious ones were involved in cholesterol and hedgehog signaling pathways (m108), as well as cadherin binding (m114).^35^

Subsequently, we wanted to investigate the effect of increased expression of the HD enriched cortical modules on clinical onset. Therefore, to investigate whether a consistent direction of effect was observed in HD patients, we assessed the direction of significant *Z*-scores for these modules in the HD clinical onset TWAS results. Mirroring the ROSMAP effects, increased expression of m108_human-cortex_ and m114 _human-cortex_ was more likely to lead to an earlier age of HD onset, while increased expression of m122_human-cortex_ and m131_human-cortex_ correlated with later age of HD onset (Fig. 6). Although only passing within-trait correction for multiple testing, modules that were also associated with other phenotypes relating to HD, included gait disturbances (e.g. m131_human-cortex_ *P*=1.20 x10^−4^). In fact, m131_human-cortex_ is the most significantly associated module with regards to gait out of all 47 ROSMAP cortical coexpression modules. We also studied whether expression patterns for the HD transcriptomic modifier genes were associated with available phenotypes in the aging cohort. These analyses revealed that *MTMR10* and *PMS2* were associated with relevant traits after correcting for multiple testing, with greatest statistical significance being achieved for working memory and global cognitive function decline (Supplementary Fig. 3, Supplementary Table 9).

Perturbagens that give rise to gene expression profiles that are similar to beneficial TWAS signatures can be prioritized for drug repurposing. We therefore selected genes from the seven HD-enriched cortical modules that displayed significant (*P*<0.05) brain-derived TWAS associations (*n*=77) and performed signature matching to over 27,000 perturbagens identified a number of highly similar (connectivity scores >95%) and dissimilar (connectivity scores <95%) perturbagens (Supplementary Table 10, Fig. 6). This gene set included three of the 15 prioritized TWAS genes (i.e. m122_human-cortex_ *PMS2* and *EIF2AK1;* as well as m2_human-cortex_ *SUMF2*). In these analyses, top perturbational classes displaying similar signatures included inhibitors of: topoisomerase, PI3K, MTOR, CDK and FLT3. Conversely, the most dissimilar compound signature was obtained for kinetin-riboside (−99.23). Top gene perturbagens with similar signatures included knockdown of the mitochondrial transporter, *SLC25A28*, as well as the NF-kappa-B inhibitor beta gene, *NFKBIB.*

**Fig. 6.**
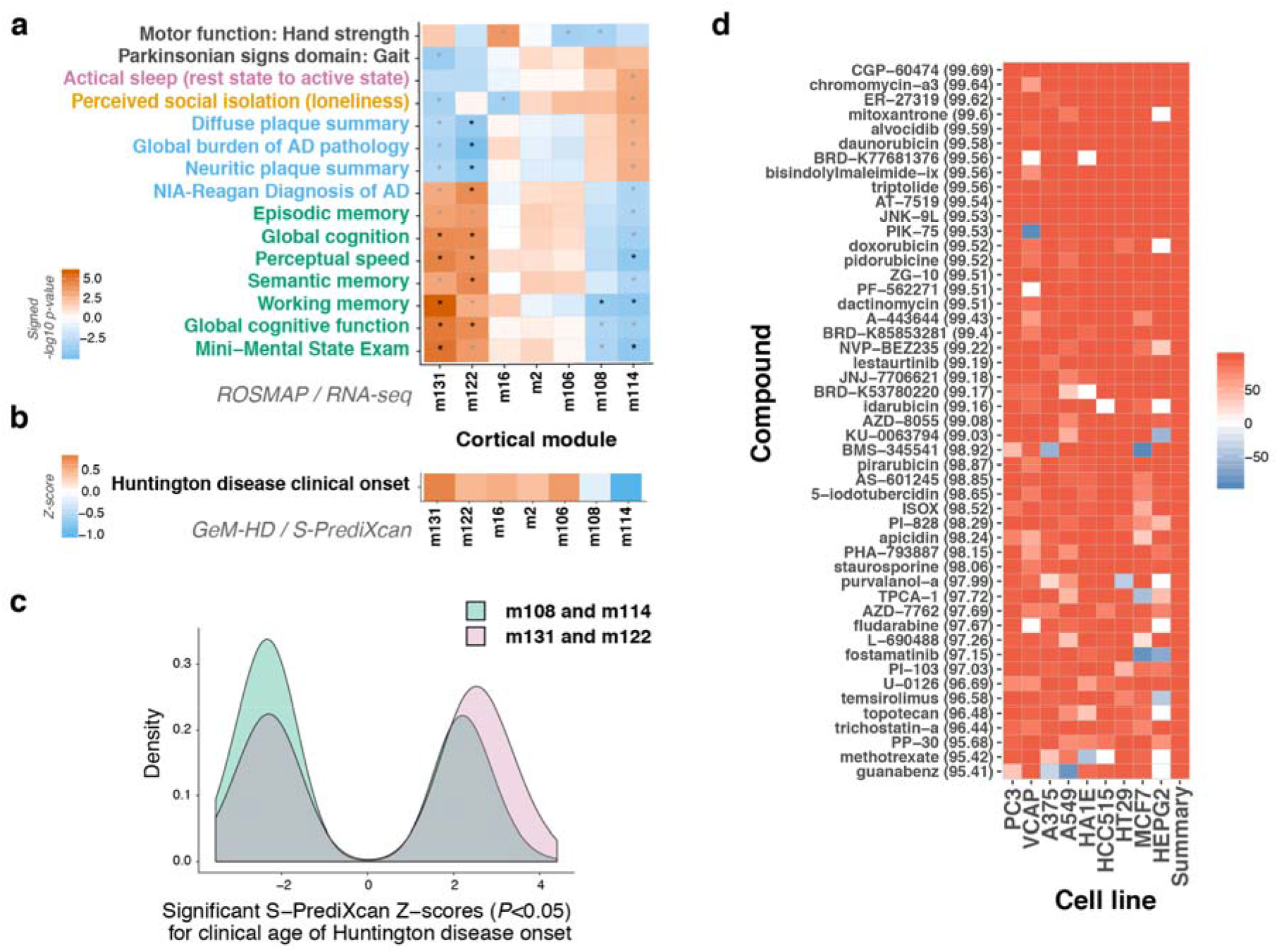
Expression of a subset of cortical gene coexpression modules influences the clinical onset of Huntington disease as well as related phenotypes. **(a)** Cortical coexpression modules that show enrichments for TWAS results for Huntington disease clinical age of onset (*n*=7) are also associated with relevant phenotypes in a longitudinal aging cohort. Increased expression of mitochondrial-related modules m122 and m131 is associated with improved performance for cognitive-related traits, while the opposite effect was observed for m108 (cholesterol and hedgehog signaling-related) and m114 (cadherin binding-related) expression. Within category Bonferroni significant traits are displayed, with asterisks denoting significant associations after Bonferroni correction for multiple testing. Trait categories are annotated (black, motor and gait; pink, sleep and circadian rhythms; orange, lifestyle/personality; blue, pathology; green, cognitive decline). Black asterisks: significant after correcting for all comparisons, grey asterisks: significant after adjusting for within trait comparisons. **(b)** A similar effect pattern was observed for these modules with regards to their influence on age of clinical onset in Huntington disease. When assessing significant S-PrediXcan *Z*-scores (i.e. *P*<0.05), on average, increased expression of genes belonging to modules m108 and m114 is more likely to lead to an earlier age of Huntington disease onset, while the opposite pattern (i.e. increased expression and later onset) is observed for the other modules. **(c)** Collectively, significant Huntington disease clinical onset S-PrediXcan *Z*-scores for the top phenotype-related coexpression cortical modules also display differences in distributions. There were significantly more negative *Z*-scores detected for genes belonging to modules m108 and m114 compared to m131 and m122 (*P*=2.0 × 10^−6^). **(d)** A screen of over 27,000 perturbagens in nine cell lines identified highly similar (≥95%) perturbagens in the Broad Connectivity Map database (median Connectivity Map / *tau* scores indicated in parentheses). Similar compounds, with potential use for drug repurposing in HD, are shown.

## DISCUSSION

Genomic approaches in HD can assist in understanding differences in clinical onset observed between patients, as well as aid in identifying novel biomarkers and therapeutic strategies to improve disease management. To our knowledge, the current study provides the most extensive TWAS of the modification of the age onset in a Mendelian disease to date, and has generated a number of novel candidate regions, as well as neurobiological pathways, for future study.

These analyses provide additional support for the role of DNA repair in disease onset, pointing towards somatic expansion of CAG tracts as one of the potential mechanisms behind this trait. By employing an integrated omic approach, we present novel evidence for another mismatch repair gene, *PMS2*, being involved in modifying the clinical onset of HD. Additionally, we show genes dysregulated in HD are enriched for disease modification signals, along with other important disease- and drug-related gene sets. Further, we identified cortical gene coexpression modules that are associated with cognitive decline as well as HD clinical onset that can also be targeted in future studies with important implications for aging populations. These findings indicate the powerful capabilities of indirectly assessing gene expression through TWAS-driven incorporation of large scale transcriptomic information into the study of genomic modifiers of disease.

Assisted by colocalization-based post-filtering, our TWAS analyses identified a number of novel modifier regions. Colocalized genes were more likely to be replicated (Table 1), forming a core list of candidates for future study. However, none of the genes in the strongest GeM-HD GWAS signal on chromosome 15q13 colocalized. This region is a complex locus, with microdeletions in this region being associated with neurodevelopmental phenotypes.^36^ The HD-related GWAS signal on this chromosome represents two independent associations and since the model assumes that there is only one causal variant in a region,^37^ the accuracy of the colocalization findings in this locus are reduced. A missense variant in the DNA repair gene*, FAN1*, could be causal for one of these signals,^38^ but the origin of the other signal at this locus still needs to be elucidated. *MTMR10* remains a good candidate for this association since it belongs to a pathway relevant to HD (i.e. inositol-phosphate signaling),^33^ as well as two trait-associated coexpression modules; a CAG-mediated huntingtin one (i.e. M7_*Htt*-striatum_)^17^ and another that forms part of the core cognitive decline related modules (i.e. m111_human-cortex_).^35^ Further, MTMR10 protein levels were increased in HD patient striatal samples (Fig. 2). This region would benefit from interrogation with functional experiments such as targeted genome editing. Other areas where such an approach should be considered include those where more than one colocalized gene is located and co-regulation is therefore suspected.

The potential role of G protein-coupled receptors (GPCRs) in modifying HD clinical onset provide an interesting line for further research. Biobank-scale genomic studies have revealed that protein-truncating *GPR151* variants are protective against obesity.^39^ Furthermore, functional characterization of neuronal *Gpr151* suggested that the gene is important in the paraventricular nucleus and could therefore be potentially implicated in food and drug seeking behaviors.^40^ Future studies should therefore aim at delineating the potential role of serotonin, glucocorticoids and these two orphan GPCRs with regards to metabolic dysfunction observed in HD, in relation to modifying age of onset of the disorder.

The TWAS also detected a number of associations with genes involved in ATP- and mitochondrial-related processes, an important finding since mitochondrial impairment has been suggested to contribute to the complex pathogenesis of HD.^3, 19^ Although previous studies in isogenic HD cell lines have provided evidence for the dysregulation of the mitochondrial-related neurodegenerative disorder risk gene, *CHCHD2*,^41^ our study provides the first evidence for the gene being involved with clinical onset in HD patients. The other TWAS gene previously implicated in neurodegenerative disorders is *EIF2AK1*^12^, although the HD signal at this locus may primarily be driven by the expression of the mismatch repair gene, *PMS2*. However, immunoblotting of EIF2AK1 in patient brains and BMECs indicated that this protein could be dysregulated in disease. Further, aberrant mRNA translation initiation and elongation have been implicated in neurodegeneration and altered in nucleotide repeat expansion disorders.^42^ The relevance of *EIF2AK1* therefore cannot be excluded.

The presence of non-brain-related ATPase-associated genes (i.e. *FXYD2* and *ATP4A*) in the prioritized gene list also warrants further investigation, but both have been implicated in processes important for HD pathogenesis, namely glutamate- and Wnt-signaling respectively.^31, 43, 44^ Like *GPR161, ATP4A* has also been shown to have important functions in the cilia,^44^ which may be relevant for HD since *HTT* is required for ciliogenesis.^19^ Finally, since sonic hedgehog signaling was implicated through both the top TWAS genes and module based analyses, further research is required in this area.

A subset of the transcriptomic modifier genes was orthogonally validated at the protein level by evaluating cortical and caudate nucleus lysates from HD patients compared to matched controls. Although not a direct comparison with regards to the age of clinical onset phenotype, we hypothesized that relevant modifier genes may also be dysregulated in the disease state. Our findings that CAG-dependent coexpression modules are enriched in TWAS associations further support this assumption. These HD clinical onset associated murine modules can be prioritized for future study to improve our biological understanding of the trait and to potentially develop therapeutic modifiers.

In addition to the *Htt*-CAG coexpression module and gene set analyses, examination of human cortical coexpression information was highly informative, indicating the mechanistic insights that can be uncovered through TWAS-driven approaches. Notably, the enriched HD-associated ROSMAP cortical modules showed shifts in their HD-onset association distributions that were correlated with their phenotypic effects in the aging individuals. The beneficial cortical modules identified here may have broader implications beyond HD, since they may be involved in processes pertaining to general brain health. The GeM-HD GWAS showed that there is a correlation between motor onset and cognitive impairment and psychiatric symptoms, and that the main HD genetic modifiers are also associated with one or more these traits.^8^ Cognitive impairments present a significant burden to the quality of life of HD patients and are often observed prior to the onset of motor symptoms.^45^ Our findings begin to provide further insights into the neurodegenerative processes that lead to cognitive deficits in HD and how they are distinct from Alzheimer’s disease. However, although ROSMAP participants had no dementia at enrolment, concomitant Alzheimer’s disease may have partially influenced these results.

Signature matching of gene expression profiles to large databases of perturbagens is a novel way to repurpose drugs to treat different diseases.^46^ Employing this approach, informed by genetic data generated from numerous HD patients, yielded a number of leads for future research. These include: chromomycin, which has been proposed as a potential therapeutic target for neurological disorders caused by repeat expansions;^47^ as well as mitoxantrone and guanabenz, which have been implicated as potential medications for multiple sclerosis.^48, 49^ Additionally, anthracyclines have been shown to correct gene expression perturbations in HD mice,^50^ while BMS-345541^51^ and the histone deacetylase inhibitor, trichostatin-a,^52^ have been shown to restore HD-related deficits *in vitro.* Further, knock down with antisense oligonucleotides of genes yielded similar expression profiles could also be explored in model systems. Conversely, kinetin-riboside, an apoptosis inducer that has been shown to cause ATP depletion and genotoxicity, ^53^ displayed the most dissimilar gene expression signature when compared to the beneficial HD modifier TWAS signals. ATP deficits are a hallmark of HD, and, while the precursor of kinetin-riboside, kinetin (also known as N6-furfuryladenine), has recently been proposed as a novel therapeutic molecule for HD,^54, 55^ kinetin-riboside may cause deleterious gene expression changes through alterations in biological feedback loops leading to DNA repair processes.

The current study is not without limitations. TWAS approaches are useful for identifying trait-related genomic regions of interest, but should not be viewed as providing causal evidence with regards to highlighted genes.^56^ Therefore, future mechanistic studies are therefore warranted for the top HD modifier genes. These include the novel modifier genes with the most supporting evidence, such as: *GPR161*, due to colocalization of signals in the cortex; *PMS2*, since the gene is a key member of the mismatch repair system; as well as the neurological disorder risk genes, *CHCHD2* and *EIF2AK1*. Additionally, although agnostic approaches to tissue selection have been recommended for TWAS,^20^ the sample size of the GTEx brain tissues is relatively limited, and future HD genomic modifier studies would benefit from the inclusion of additional striatal and cortical samples. A recent, more targeted HD modifier study compared to the current investigation, used dorsolateral prefrontal cortex eQTL information (*n*=452) from the CommonMind Consortium only provided additional evidence for the *FAN1* gene.^57^ Although the HD patients in the replication study were extensively phenotyped for disease progression,^7^ the sample size of this cohort also limited our power to confirm associations. Finally, the applicability of TWAS are inherently limited with regards to the importance of steady-state gene expression on the phenotype being studied. Our gene expression informed analyses identified a number of novel findings, but further in-depth analyses of the role of deleterious coding variation is also required.

In conclusion, by jointly modelling HD clinical onset GWAS summary statistics with population-scale tissue-specific eQTL information, we expanded the number of candidate modifier genes and gained increased insights into HD disease mechanisms. Transcriptomic imputation can therefore be seen as a complementary approach to GWAS, as it can reveal novel information not previously detected with traditional methodologies. Significant progress can be made if these findings help inform clinical translation by paving the way forward for comprehensive drug repurposing. Further, expanding future TWAS to incorporate modifiers of other neurodegenerative disorders will aid in identifying common pathways across these diseases, as well to ascertain those that are unique to each disorder.

## METHODS

### Transcriptome-wide association study and Bayesian colocalization analyses

S-PrediXcan^20^ was used to perform a TWAS by integrating gene expression prediction models (http://predictdb.org/) generated from the GTEx Consortium (v7 release)^33^ with summary statistics from two GWAS of the genetic modifiers of HD. These models were built using eQTL information from 48 tissues, which includes 13 brain regions. GWAS data from 4,082 European HD patients (CAG ranges: 40-55) generated by the Genetic Modifiers of Huntington’s Disease (GeM-HD) Consortium^8^ was used in the discovery analysis. Analyses were restricted to protein coding genes and a false discovery rate (FDR) of 0.1 for each tissue was used to determine statistical significance. Since eQTL data can display correlation across tissues,^33^ we selected this threshold in an attempt to balance minimizing detecting false-positive associations, while maximizing discovery power.

To validate prioritized genes further, filtering was performed employing Bayesian colocalization analyses with COLOC.^37^ Non-ambiguous variants in the GeM-HD GWAS data were interrogated in relation to the eQTL association results from the GTEx Consortium,^33^ as well as those generated from dorsolateral prefrontal cortex samples from 411 research participants as part of a longitudinal study of aging. COLOC produces posterior probabilities for four hypotheses: *H0, no causal variant; H1, causal variant for trait 1 only; H2, causal variant for trait 2 only; H3, two distinct causal variants; H4, one common causal variant.* In order to be defined as colocalized in a tissue, regions were required to show strong evidence for one common causal variant (i.e. P4>0.5), without substantial evidence for the PrediXcan associations occurring as a result of linkage disequilibrium contamination.

Regions that were significantly associated in the discovery analysis and were validated through Bayesian colocalization analyses were further examined for replication. We assessed replication in S-PrediXcan gene level associations generated using GWAS of disease progression in 216 European HD patients recruited by the TRACK-HD Consortium.^7^ We considered a gene to be replicated at a nominal *P*<0.05 when the association effect was in the same direction in significant discovery and replication tissues.

The biomaRt package in R was used to access Ensembl database (grch37.ensembl.org) to obtain relevant genomic coordinates of genes of interest as well as mouse gene homologues. Networks of physical and other interactions based on diverse genomic and proteomic data with gene sets of interest were visualized using GeneMANIA.^58^

### Orthogonal validation of prioritized transcriptomic modifier genes using data from HD allelic hPSCs, HD knock-in mice and human HD patient brains

Gene expression of prioritized genes was assessed using RNA-seq data from isogenic allelic hPSC lines for four different *HTT* CAG lengths (CAG lengths: 30, 45, 65, 81; *n*=4 per length) that had been differentiated into five cell line types: pluripotent stem cells, hepatocytes, neural progenitor cells, post-mitotic neurons and skeletal muscle cells. Further details are described in Ooi *et al.^59^* The expression for each gene (transcripts per kilobase million) and cell line combination were assessed as a continuous trait with linear regression for different CAG lengths.

HD knock-in mouse RNA-seq data from Langfelder *et al.*^17^ were used to determine whether prioritized transcriptomic HD modifier gene were differentially expressed Q175 HD mice (*n*=8) versus Q20 wild type mice (*n*=8) at 6 months for 12 tissues, including six brain-related regions.

Protein levels for a subset of the prioritized genes (i.e. *CHCHD2, EIF2AK1, FAN1, GPR161, MSH3, MTMR10, PMS2, RRM2B* and *SUMF2)* were assessed using Western blot analysis in HD patient brains (*n*=10) compared to age-matched controls (*n*=9) in both the striatum (caudate nucleus) and cortex. These samples were obtained from deidentified archived brain tissue samples from the Huntington Disease Biobank at the University of British Columbia. All samples were collected, stored and accessed with informed consent and approval of the University of British Columbia / Children’s and Women’s Health Centre of British Columbia Research Ethics Board (UBC C&W REB H06-70467 and H06-70410)

For immunoblotting, a small piece (2mm × 2mm) of frozen cortex or caudate nucleus was cut using a clean razor blade on dry ice. Each brain section was lysed and 75 μg of total protein was resolved on a 12% SDS-PAGE gel. Proteins were transferred to 0.2μm nitrocellulose membranes that were blotted for either GPR161 (1:250, Abcam, ab 83007), EIF2AK1 (1:500, Abcam, ab154076), PMS2 (1:500, Abcam, ab110638), MTMR10 (1:250, Abcam, ab179756), CHCHD2 (1:500, Abcam, ab220688), MSH3 (1:250, Abcam, ab154486), FAN1 (1:1000, Abcam, ab185554), SUMF2 (1:250, ThermoFisher, PA5-54961), RRM2B (p53R2, 1:1000, Abcam, ab154194) and actin (1:5000, Sigma-Aldrich, A2228). Proteins were detected with IR dye 800CW goat antimouse (1:250, Rockland 610-131-007, Gilbertsville, PA) and AlexaFluor 680 goat antirabbit (1:250, Molecular Probes A21076, Eugene, OR)-labeled secondary antibodies and the LiCor Odyssey Infrared Imaging system.

### Assessing important cell types for HD clinical onset using single cell sequencing information

Relevant cell types for modifying HD clinical onset were inferred using RolyPoly,^60^ which employs a regression-based polygenic model to detect enrichment of GWAS signal based on gene expression information for different cell populations. Developer recommended criteria were implemented using publically-available single cell sequencing data from human cortex.^61^ These data had been clustered into six major brain cell types: astrocytes, oligodendrocytes, oligodendrocyte precursor cells, neurons, microglia, and vascular cells, as well as replicating and quiescent fetal neuronal populations.^61^ Significant differences in expression between HD patients and controls for prioritized transcriptomic HD modifier genes in brain microvascular endothelial cells was assessed using RNA-seq information from a previous study, as described by the authors.^31^

### Brain microvascular endothelial cell differentiation

GM09197 HD, C1 and C2 induced hPSC lines were generated as described in Xu *et al.^41^* GM03621 HD fibroblasts were obtained from the NIGMS Human Genetic Cell Repository at the Coriell Institute for Medical Research. Induced hPSCs from this line were generated using an episomal reprogramming strategy as described in Okita *et al.*^62^ and characterized for pluripotency and karyotype. Differentiation to brain microvascular endothelial cells was performed using a protocol modified from Lippmann *et al.*^63^ Briefly, iPSCs were grown in mTeSR1 (STEMCELL Technology cat# 85850) and were passaged as aggregates onto Matrigel coated plates (Corning cat# 354230). Cells were grown to 40% confluency before changing media to DMEM/F12 (Gibco cat# 11320033) supplemented 20% KnockOut serum replacement (Gibco cat # 10828028), 1 × MEM Non-Essential Amino Acids Solution (Gibco cat# 11140050), 1 × GlutaMAX-I (Gibco cat # 35050061) and 0.1mM beta-mercaptoethanol. DMEM/F12 complete media was changed daily for 5 days. On day 6, media was changed to human endothelial media (Gibco cat# 11111044) supplemented with 20ng/mL bFGF (ProSpec cat # CYT-218) and 1% platelet poor-derived serum (Sigma cat# P2918). Endothelial media was changed daily for 2 days. On day 8, cells were dissociated with accutase and seeded at 25,000 cells/cm2 on collagen IV (400ug/mL; Sigma C5533) and fibronectin (100ug/mL; Sigma F4759) coated plates. Endothelial media was changed daily for four days and confluent cells were harvested for immunoblotting.

Enrichment of transcriptomic gene level association results for HD onset in gene sets We performed TWAS association enrichment analyses (mean S-PrediXcan *Z*^2^-score for all gene-tissue pairs, as previously described^20^) with regards to diverse gene sets in the GeM-HD cohort. Bonferroni correction for multiple testing was applied to determine statistically significant enrichment of gene sets. Seventeen gene sets (https://github.com/macarthur-lab/gene_lists; downloaded November 2017, *n*=17) from a repository which includes annotations for genes grouped according to biological function, involvement in specific inheritance models of diseases and known drug targets (complete descriptions can be found in Supplementary Table 4), were assessed. Further, gene sets belonging to coexpression modules identified through RNA-seq analysis in brain regions most relevant to HD (i.e. cortex and striatum)^3^ were also assessed for enrichment in the TWAS data. These gene sets included striatal and cortical coexpression modules that are mediated by murine *Htt*-CAG length and age (six CAG lengths and three time points, *n=144* mice per tissue)^17^ and coexpression modules identified in the human cortex (*n=*478)^35^.

### Relevance of genetic modifiers of HD age of onset to related traits in the aging brain

The human cortical samples (*n*=478) used to identify coexpression modules relevant to modifying HD clinical onset were collected from two prospective cohorts (ROSMAP), which extensively characterized the study participants for cognitive measures for up to 20 years and performed pathological characterization post-mortem (https://www.radc.rush.edu/documentation.htm).^35^ This allowed for the opportunity to determine the relevance of the genetic HD modifiers to related phenotypes observed in the aging brain. Cortical coexpression modules (*n*=47) that displayed significant enrichments for S-PrediXcan associations with HD clinical age of onset were examined with regards to clinical phenotypes in a longitudinal aging cohort. In these analyses, the expression of significantly enriched HD modifier modules (*n*=7) were assessed for association with clinical and pathological traits (*n*=83), including those relevant to HD, in the ROSMAP cohort. Modules that displayed significant associations with at least one ROSMAP trait were examined with regards to the direction of significant *Z*-scores (*P*<0.05) in the HD clinical age of onset TWAS. Cortically-expressed prioritized HD transcriptomic modifier genes, as well as *MSH3* and *HTT*, were also assessed with regards to associations with these ROSMAP phenotypes. Individual gene expression levels were only available for a subset of the phenotypes present in the ROSMAP cohort examined in Mostafavi *et al*.^35^ (*n*=13), thus gene level analyses were performed in this subset of traits.

Association between each module and each trait was conducted by first representing each module by the mean gene expression level of the module’s gene members (applied to standardized gene expression data), and then using Spearman correlation to associate module means with observed phenotypic traits. Bonferroni correction for multiple testing was used to determine statistical significance with traits and genes (significance thresholds: module-trait analyses *P*<8.6 × 10^−5^ and gene-multi-trait analyses *P*<3.2 × 10^−4^. Nominal significance for gene-individual-trait analyses *P*<0.004).

### Perturbagen signature matching for potential drug repurposing in Huntington disease

HD modifier TWAS signals matched to gene expression profiles from over 27,000 perturbagens in nine cell lines in ConnectivityMap (CMap) database^64^ (https://clue.io/). TWAS *Z*-scores were used as a proxy for gene expression (i.e. positive *Z*-scores as upregulated genes and negative *Z*-scores as down regulated ones), with perturbagens giving rise to highly similar median CMap connectivity score profiles (i.e. >95%) being considered potential targets for drug repurposing. These analyses were restricted to genes that fulfilled all of the following criteria: (i) members of one of the seven cortical modules from ROSMAP that displayed enrichment for HD onset associations, (ii) displaying at least suggestive TWAS significance (*P*<0.05) in at least one GTEx brain tissue, and (iii) the direction of these TWAS *Z*-scores should match the direction of the predicted mean influence on HD onset of the co-expression module (i.e. positive TWAS *Z*-scores for m2, m16, m106, m122 and m131, and negative TWAS *Z*-scores for m108 and m114).

All downstream statistical analyses were conducted using R version 3.4.4 software (The R Foundation).

## ACKNOWLEDGEMENTS

We would like to acknowledge the GeM-HD and TRACK-HD Consortia for providing access to their GWAS summary statistics. This work was supported by a Canadian Institutes of Health Research Foundation Grant awarded to M.R.H.

## REFERENCES

1. Harper, A.R., Nayee, S. & Topol, E.J. Protective alleles and modifier variants in human health and disease. Nat Rev Genet 16, 689–701 (2015).

2. Caron, N.S., Wright, G.E.B. & Hayden, M.R. Huntington Disease, in GeneReviews((R)) (ed. M.P. Adam, et al.) (Seattle (WA), 2018).

3. Bates, G.P., et al. Huntington disease. Nat Rev Dis Primers 1, 15005 (2015).

4. Keum, J.W., et al. The HTT CAG-Expansion Mutation Determines Age at Death but Not Disease Duration in Huntington Disease. American Journal of Human Genetics 98, 287–298 (2016).

5. Gusella, J.F. & MacDonald, M.E. Huntington’s disease: the case for genetic modifiers. Genome Med 1, 80 (2009).

6. Rosenblatt, A., et al. Familial influence on age of onset among siblings with Huntington disease. Am J Med Genet 105, 399–403 (2001).

7. Hensman Moss, D.J., et al. Identification of genetic variants associated with Huntington’s disease progression: a genome-wide association study. Lancet Neurol 16, 701–711 (2017).

8. GeM-HD Consortium. Identification of Genetic Factors that Modify Clinical Onset of Huntington’s Disease. Cell 162, 516–526 (2015).

9. Becanovic, K., et al. A SNP in the HTT promoter alters NF-kappaB binding and is a bidirectional genetic modifier of Huntington disease. Nat Neurosci 18, 807–816 (2015).

10. GeM-HD Consortium, et al. Huntington’s disease onset is determined by length of uninterrupted CAG, not encoded polyglutamine, and is modified by DNA maintenance mechanisms. bioRxiv, 529768 (2019).

11. Wright, G.E.B., et al. Length of Uninterrupted CAG, Independent of Polyglutamine Size, Results in Increased Somatic Instability, Hastening Onset of Huntington Disease. American Journal of Human Genetics (2019).

12. Sadovnick, A.D., et al. Genetic modifiers of multiple sclerosis progression, severity and onset. Clin Immunol 180, 100–105 (2017).

13. van Blitterswijk, M., et al. TMEM106B protects C9ORF72 expansion carriers against frontotemporal dementia. Acta Neuropathol 127, 397–406 (2014).

14. Deming, Y., et al. Genome-wide association study identifies four novel loci associated with Alzheimer’s endophenotypes and disease modifiers. Acta Neuropathol 133, 839–856 (2017).

15. Bettencourt, C., et al. DNA repair pathways underlie a common genetic mechanism modulating onset in polyglutamine diseases. Ann Neurol 79, 983–990 (2016).

16. Hauberg, M.E., et al. Large-Scale Identification of Common Trait and Disease Variants Affecting Gene Expression. American Journal of Human Genetics 100, 885–894 (2017).

17. Langfelder, P., et al. Integrated genomics and proteomics define huntingtin CAG length-dependent networks in mice. Nat Neurosci 19, 623–633 (2016).

18. Ament, S.A., et al. Transcriptional regulatory networks underlying gene expression changes in Huntington’s disease. Mol Syst Biol 14, e7435 (2018).

19. Saudou, F. & Humbert, S. The Biology of Huntingtin. Neuron 89, 910–926 (2016).

20. Barbeira, A.N., et al. Exploring the phenotypic consequences of tissue specific gene expression variation inferred from GWAS summary statistics. Nat Commun 9, 1825 (2018).

21. Gusev, A., et al. Integrative approaches for large-scale transcriptome-wide association studies. Nature genetics 48, 245–252 (2016).

22. Zhu, Z., et al. Integration of summary data from GWAS and eQTL studies predicts complex trait gene targets. Nature Genetics 48, 481–487 (2016).

23. Ng, B., et al. An xQTL map integrates the genetic architecture of the human brain’s transcriptome and epigenome. Nat Neurosci 20, 1418–1426 (2017).

24. ten Broeke, S.W., et al. Lynch syndrome caused by germline PMS2 mutations: delineating the cancer risk. Journal of Clinical Oncology 33, 319–325 (2015).

25. Ogaki, K., et al. Mitochondrial targeting sequence variants of the CHCHD2 gene are a risk for Lewy body disorders. Neurology 85, 2016–2025 (2015).

26. Hormozdiari, F., et al. Colocalization of GWAS and eQTL Signals Detects Target Genes. American Journal of Human Genetics 99, 1245–1260 (2016).

27. Shimada, I.S., et al. Basal Suppression of the Sonic Hedgehog Pathway by the G-Protein-Coupled Receptor Gpri6i Restricts Medulloblastoma Pathogenesis. Cell Rep 22, 1169–1184 (2018).

28. Aravind, L., Walker, D.R. & Koonin, E.V. Conserved domains in DNA repair proteins and evolution of repair systems. Nucleic Acids Res 27, 1223–1242 (1999).

29. Steinberg, S., et al. Loss-of-function variants in ABCA7 confer risk of Alzheimer’s disease. Nature Genetics 47, 445–447 (2015).

30. Wiatr, K., Szlachcic, W.J., Trzeciak, M., Figlerowicz, M. & Figiel, M. Huntington Disease as a Neurodevelopmental Disorder and Early Signs of the Disease in Stem Cells. Mol Neurobiol 55, 3351–3371 (2018).

31. Lim, R.G., et al. Huntington’s Disease iPSC-Derived Brain Microvascular Endothelial Cells Reveal WNT-Mediated Angiogenic and Blood-Brain Barrier Deficits. Cell Rep 19, 1365–1377 (2017).

32. Landrum, M.J., et al. ClinVar: public archive of relationships among sequence variation and human phenotype. Nucleic Acids Res 42, D980–985 (2014).

33. Consortium, G.T., et al. Genetic effects on gene expression across human tissues. Nature 550, 204–213 (2017).

34. Nelson, M.R., et al. An abundance of rare functional variants in 202 drug target genes sequenced in 14,002 people. Science 337, 100–104 (2012).

35. Mostafavi, S., et al. A molecular network of the aging human brain provides insights into the pathology and cognitive decline of Alzheimer’s disease. Nat Neurosci 21, 811–819 (2018).

36. Uddin, M., et al. OTUD7A Regulates Neurodevelopmental Phenotypes in the 15q13.3 Microdeletion Syndrome. American Journal of Human Genetics 102, 278–295 (2018).

37. Giambartolomei, C., et al. Bayesian test for colocalisation between pairs of genetic association studies using summary statistics. PLoS Genet 10, e1004383 (2014).

38. Chao, M.J., et al. Population-specific genetic modification of Huntington’s disease in Venezuela. PLoS Genet 14, e1007274 (2018).

39. Emdin, C.A., et al. Analysis of predicted loss-of-function variants in UK Biobank identifies variants protective for disease. Nat Commun 9, 1613 (2018).

40. Broms, J., et al. Monosynaptic retrograde tracing of neurons expressing the G-protein coupled receptor Gpr151 in the mouse brain. J Comp Neurol 525, 3227–3250 (2017).

41. Xu, X., et al. Reversal of Phenotypic Abnormalities by CRISPR/Cas9-Mediated Gene Correction in Huntington Disease Patient-Derived Induced Pluripotent Stem Cells. Stem Cell Reports 8, 619–633 (2017).

42. Kapur, M., Monaghan, C.E. & Ackerman, S.L. Regulation of mRNA Translation in Neurons-A Matter of Life and Death. Neuron 96, 616–637 (2017).

43. Kinoshita, P.F., et al. The Influence of Na(+), K(+)-ATPase on Glutamate Signaling in Neurodegenerative Diseases and Senescence. Front Physiol 7, 195 (2016).

44. Walentek, P., Beyer, T., Thumberger, T., Schweickert, A. & Blum, M. ATP4a is required for Wnt-dependent Foxj_1_ expression and leftward flow in Xenopus left-right development. Cell Rep 1, 516–527 (2012).

45. Paulsen, J.S., Smith, M.M., Long, J.D., investigators, P.H. & Coordinators of the Huntington Study, G. Cognitive decline in prodromal Huntington Disease: implications for clinical trials. J Neurol Neurosurg Psychiatry 84, 1233–1239 (2013).

46. Pushpakom, S., et al. Drug repurposing: progress, challenges and recommendations. Nat Rev Drug Discov (2018).

47. Chen, Y.W., et al. Co(II)(Chromomycin)(2) Complex Induces a Conformational Change of CCG Repeats from i-Motif to Base-Extruded DNA Duplex. Int J Mol Sci 19 (2018).

48. Martinelli Boneschi, F., Vacchi, L., Rovaris, M., Capra, R. & Comi, G. Mitoxantrone for multiple sclerosis. Cochrane Database Syst Rev, CD002127 (2013).

49. Way, S.W., et al. Pharmaceutical integrated stress response enhancement protects oligodendrocytes and provides a potential multiple sclerosis therapeutic. Nat Commun 6, 6532 (2015).

50. Stack, E.C., et al. Modulation of nucleosome dynamics in Huntington’s disease. Hum Mol Genet 16, 1164–1175 (2007).

51. Caron, N.S., Desmond, C.R., Xia, J. & Truant, R. Polyglutamine domain flexibility mediates the proximity between flanking sequences in huntingtin. Proc Natl Acad Sci USA 110, 14610–14615 (2013).

52. Naia, L., et al. Histone Deacetylase Inhibitors Protect Against Pyruvate Dehydrogenase Dysfunction in Huntington’s Disease. J Neurosci 37, 2776–2794 (2017).

53. Cabello, C.M., et al. The experimental chemotherapeutic N6-furfuryladenosine (kinetinriboside) induces rapid ATP depletion, genotoxic stress, and CDKN1A(p21) upregulation in human cancer cell lines. Biochem Pharmacol 77, 1125–1138 (2009).

54. Bowie, L.E., et al. N6-Furfuryladenine is protective in Huntington’s disease models by signaling huntingtin phosphorylation. Proc Natl Acad Sci USA 115, E7081–E7090 (2018).

55. Maiuri, T., Bowie, L.E. & Truant, R. DNA Repair Signaling of Huntingtin: The Next Link Between Late-Onset Neurodegenerative Disease and Oxidative DNA Damage. DNA Cell Biol 38, 1–6 (2019).

56. Wainberg, M., et al. Opportunities and challenges for transcriptome-wide association studies. Nature Genetics 51, 592–599 (2019).

57. Goold, R., et al. FAN_1_ modifies Huntington’s disease progression by stabilising the expanded HTT CAG repeat. Hum Mol Genet (2018).

58. Franz, M., et al. GeneMANIA update 2018. Nucleic Acids Res 46, W60–W64 (2018).

59. Ooi, J., et al. Unbiased Profiling of Isogenic Huntington Disease hPSC-Derived CNS and Peripheral Cells Reveals Strong Cell-Type Specificity of CAG Length Effects. Cell Rep 26, 2494–2508 e2497 (2019).

60. Calderon, D., et al. Inferring Relevant Cell Types for Complex Traits by Using Single-Cell Gene Expression. American Journal of Human Genetics 101, 686–699 (2017).

61. Darmanis, S., et al. A survey of human brain transcriptome diversity at the single cell level. Proc Natl Acad Sci USA 112, 7285–7290 (2015).

62. Okita, K., et al. A more efficient method to generate integration-free human iPS cells. Nat Methods 8, 409–412 (2011).

63. Lippmann, E.S., et al. Derivation of blood-brain barrier endothelial cells from human pluripotent stem cells. Nat Biotechnol 30, 783–791 (2012).

64. Subramanian, A., et al. A Next Generation Connectivity Map: L1000 Platform and the First 1,000,000 Profiles. Cell 171, 1437–1452 e1417 (2017).

