## Supplementary Tables S1-S10 and Figures S1-S4 for "Gene expression profiles complement the analysis of genomic modifiers of the clinical onset of Huntington disease"

### SUPPLEMENTARY INFORMATION

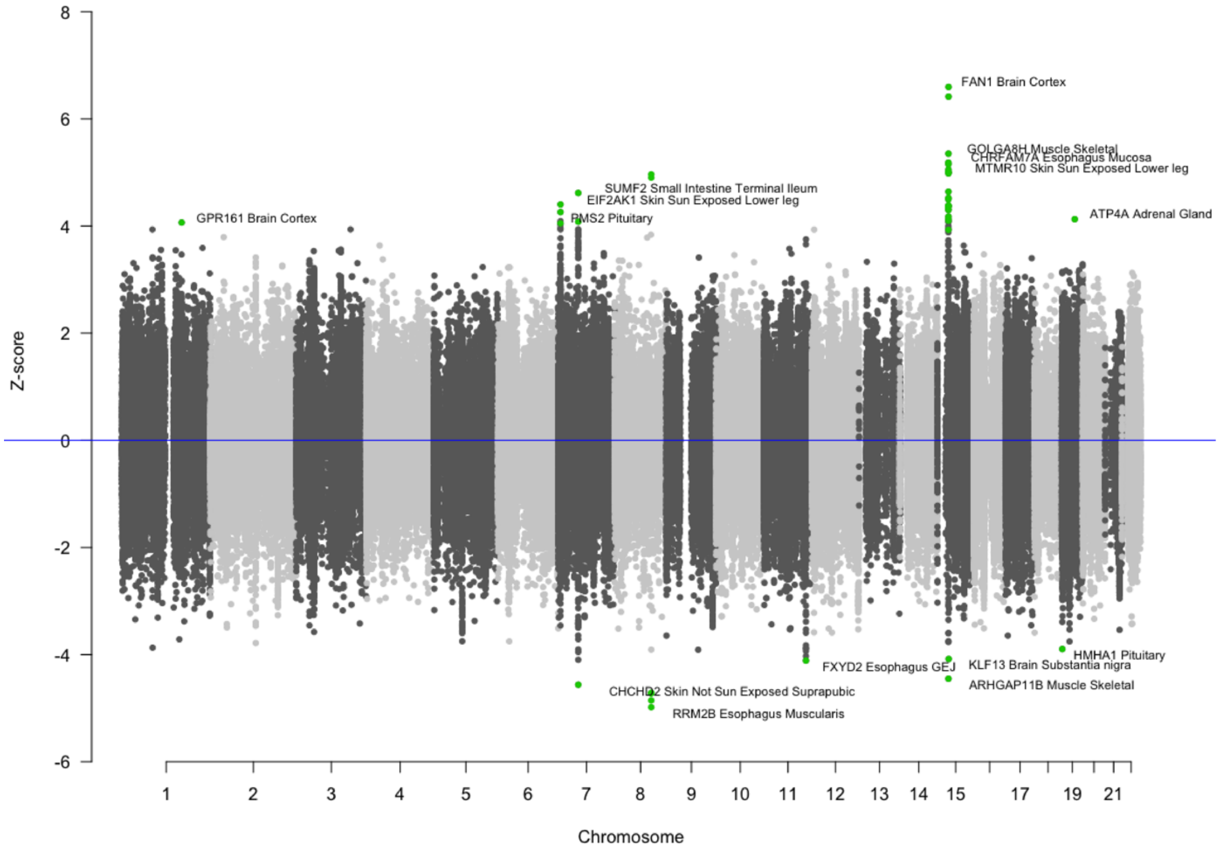


**Supplementary Fig. 1** Miami plot displaying the association results from the HD clinical onset TWAS. S-PrediXcan *Z*-scores are indicated on the *y*-axis and chromosomal location of genes on the *x*-axis. The most significant gene-tissue association for each of the 15 prioritized transcriptomic modifier genes are annotated in green.

**Supplementary Table 1** Gene-tissue pairs for the genetic modifiers of the clinical onset of Huntington disease that passed the false discovery rate of 0.1 correction for multiple testing.

| Tissue | Gene | GeM-HD *Z*-score | GeM-HD *P*-value |
|---|---|---|---|
| Brain Cortex | *FAN1* | 6.60 | 4.25E-11 |
| Adipose Subcutaneous | *FAN1* | 6.42 | 1.41E-10 |
| Muscle Skeletal | *GOLGA8H* | 5.35 | 8.73E-08 |
| Esophagus Mucosa | *CHRFAM7A* | 5.18 | 2.19E-07 |
| Heart Left Ventricle | *GOLGA8H* | 5.15 | 2.55E-07 |
| Nerve Tibial | *GOLGA8H* | 5.05 | 4.42E-07 |
| Breast Mammary Tissue | *GOLGA8H* | 5.01 | 5.57E-07 |
| Skin Sun Exposed Lower leg | *MTMR10* | 5.00 | 5.79E-07 |
| Artery Tibial | *GOLGA8H* | 4.98 | 6.28E-07 |
| Esophagus Muscularis | *RRM2B* | -4.98 | 6.29E-07 |
| Esophagus Mucosa | *RRM2B* | 4.96 | 6.91E-07 |
| Muscle Skeletal | *RRM2B* | 4.90 | 9.52E-07 |
| Thyroid | *RRM2B* | -4.86 | 1.20E-06 |
| Artery Tibial | *RRM2B* | -4.72 | 2.32E-06 |
| Esophagus Gastroesophageal Junction | *RRM2B* | -4.71 | 2.44E-06 |
| Adrenal Gland | *GOLGA8H* | 4.64 | 3.46E-06 |
| Small Intestine Terminal Ileum | *SUMF2* | 4.62 | 3.89E-06 |
| Skin Not Sun Exposed Suprapubic | *CHCHD2* | -4.56 | 5.10E-06 |
| Esophagus Muscularis | *GOLGA8H* | 4.52 | 6.08E-06 |
| Artery Coronary | *CHRFAM7A* | 4.50 | 6.82E-06 |
| Muscle Skeletal | *ARHGAP11B* | -4.45 | 8.63E-06 |
| Skin Sun Exposed Lower leg | *EIF2AK1* | 4.40 | 1.07E-05 |
| Pituitary | *GOLGA8H* | 4.38 | 1.16E-05 |
| Artery Coronary | *GOLGA8H* | 4.37 | 1.22E-05 |
| Skin Not Sun Exposed Suprapubic | *GOLGA8H* | 4.37 | 1.26E-05 |

| | | | |
|---|---|---|---|
| Esophagus Gastroesophageal Junction | *CHRFAM7A* | 4.37 | 1.27E-05 |
| Brain Putamen basal ganglia | *CHRFAM7A* | 4.32 | 1.58E-05 |
| Adipose Visceral Omentum | *GOLGA8H* | 4.30 | 1.73E-05 |
| Heart Atrial Appendage | *EIF2AK1* | 4.26 | 2.04E-05 |
| Adipose Subcutaneous | *GOLGA8H* | 4.18 | 2.94E-05 |
| Skin Sun Exposed Lower leg | *GOLGA8H* | 4.13 | 3.57E-05 |
| Adrenal Gland | *ATP4A* | 4.13 | 3.67E-05 |
| Esophagus Gastroesophageal Junction | *FXYD2* | -4.11 | 3.94E-05 |
| Ovary | *GOLGA8H* | 4.10 | 4.17E-05 |
| Pituitary | *SUMF2* | 4.08 | 4.50E-05 |
| Brain Substantia nigra | *KLF13* | -4.08 | 4.50E-05 |
| Brain Cortex | *GPR161* | 4.06 | 4.82E-05 |
| Pituitary | *PMS2* | 4.05 | 5.09E-05 |
| Brain Cortex | *GOLGA8H* | 3.93 | 8.49E-05 |
| Pituitary | *HMHA1* | -3.90 | 9.78E-05 |

**Supplementary Table 2** Top transcriptomic modifier HD genes that also displayed replication in the TRACK-HD cohort with effects in the same direction across cohorts.

| Tissue | Gene | GeM-HD *Z*-score | GeM-HD *P*-value | TRACK-HD *Z*-score | TRACK-HD *P*-value |
|---|---|---|---|---|---|
| Heart Atrial Appendage | *EIF2AK1* | 4.26 | 2.04E-05 | 2.68 | 7.33E-03 |
| Cells EBV-transformed lymphocytes | *EIF2AK1* | 4.09 | 4.30E-05 | 2.89 | 3.81E-03 |
| Pituitary | *SUMF2* | 4.08 | 4.50E-05 | 2.18 | 2.90E-02 |
| Skin Not Sun Exposed Suprapubic | *EIF2AK1* | 4.03 | 5.69E-05 | 2.22 | 2.61E-02 |
| Adipose Subcutaneous | *CHCHD2* | -3.50 | 4.74E-04 | -2.12 | 3.43E-02 |
| Skin Sun Exposed Lower leg | *CHCHD2* | -3.38 | 7.38E-04 | -2.29 | 2.19E-02 |
| Uterus | *CHCHD2* | -3.29 | 9.91E-04 | -2.13 | 3.33E-02 |
| Adrenal Gland | *PMS2* | 3.28 | 1.04E-03 | 2.87 | 4.13E-03 |
| Artery Coronary | *PMS2* | 3.28 | 1.04E-03 | 2.13 | 3.33E-02 |
| Adrenal Gland | *EIF2AK1* | 3.21 | 1.31E-03 | 2.69 | 7.19E-03 |
| Cells Transformed fibroblasts | *EIF2AK1* | 3.17 | 1.52E-03 | 2.00 | 4.53E-02 |
| Adipose Visceral Omentum | *PMS2* | 3.03 | 2.48E-03 | 2.23 | 2.55E-02 |
| Artery Aorta | *EIF2AK1* | 2.91 | 3.59E-03 | 3.23 | 1.25E-03 |
| Adipose Subcutaneous | *EIF2AK1* | 2.90 | 3.77E-03 | 2.60 | 9.43E-03 |
| Adipose Visceral Omentum | *EIF2AK1* | 2.88 | 4.03E-03 | 3.02 | 2.55E-03 |
| Heart Atrial Appendage | *PMS2* | 2.73 | 6.36E-03 | 2.06 | 3.93E-02 |
| Testis | *EIF2AK1* | 2.58 | 9.99E-03 | 2.19 | 2.88E-02 |
| Whole Blood | *CHCHD2* | -2.40 | 1.64E-02 | -2.03 | 4.19E-02 |
| Esophagus Mucosa | *MTMR10* | -2.39 | 1.68E-02 | -2.20 | 2.75E-02 |
| Pancreas | *PMS2* | 2.38 | 1.75E-02 | 2.02 | 4.38E-02 |
| Cells Transformed fibroblasts | *PMS2* | 2.35 | 1.89E-02 | 2.57 | 1.02E-02 |
| Heart Atrial Appendage | *MTMR10* | -2.28 | 2.24E-02 | -1.97 | 4.86E-02 |
| Whole Blood | *PMS2* | 1.96 | 4.96E-02 | 2.05 | 4.04E-02 |

**Supplementary Table 3** Differences in protein levels of top HD modifiers ascertained by Western blotting in HD patient and age-matched control brain tissues.

| Protein | Tissue | *P*-value | Mean controls (SEM) | Mean HD (SEM) |
|---|---|---|---|---|
| *CHCHD2* | Caudate nucleus | 0.9717 | 1.000 ± 0.2874 N=5 | 0.9855 ± 0.2720 N=5 |
| *EIF2AK1* | Caudate nucleus | 0.0015 | 1.000 ± 0.1441 N=9 | 0.3635 ± 0.08275 N=9 |
| *FAN1* (67 KDa) | Caudate nucleus | 0.1974 | 1.000 ± 0.09031 N=5 | 0.7648 ± 0.1408 N=5 |
| *FAN1* (150 KDa) | Caudate nucleus | 0.0537 | 1.000 ± 0.2186 N=5 | 0.3878 ± 0.1598 N=5 |
| *GPR161* | Caudate nucleus | 0.005 | 1.000 ± 0.1061 N=5 | 0.4514 ± 0.09592 N=5 |
| *MSH3* | Caudate nucleus | 0.0399 | 1.000 ± 0.1758 N=10 | 0.5389 ± 0.06528 N=8 |
| *MTMR10* | Caudate nucleus | 0.0098 | 1.000 ± 0.1557 N=9 | 2.027 ± 0.4595 N=8 |
| *RRM2B* | Caudate nucleus | 0.2076 | 1.000 ± 0.1634 N=10 | 0.7273 ± 0.1230 N=9 |
| *SUMF2* | Caudate nucleus | 0.1861 | 1.000 ± 0.1025 N=10 | 0.8275 ± 0.06616 N=9 |
| *PMS2* | Caudate nucleus | 0.1104 | 1.000 ± 0.3046 N=5 | 0.3972 ± 0.1413 N=5 |
| *CHCHD2* | Cortex | 0.3069 | 1.000 ± 0.2875 N=5 | 1.517 ± 0.3763 N=5 |
| *EIF2AK1* | Cortex | 0.8529 | 1.000 ± 0.1051 N=8 | 0.9685 ± 0.1294 N=8 |
| *FAN1* (67 KDa) | Cortex | 0.0004 | 1.000 ± 0.06128 N=5 | 0.5207 ± 0.03467 N=4 |
| *FAN1* (150 KDa) | Cortex | 0.0491 | 1.000 ± 0.1111 N=5 | 0.6642 ± 0.07271 N=4 |
| *GPR161* | Cortex | 0.1359 | 1.000 ± 0.1401 N=5 | 0.6592 ± 0.1426 N=4 |
| *MSH3* | Cortex | 0.9874 | 1.000 ± 0.1928 N=8 | 0.9952 ± 0.2292 N=7 |
| *MTMR10* | Cortex | 0.0071 | 1.000 ± 0.08882 N=8 | 0.6145 ± 0.08408 N=8 |
| *RRM2B* | Cortex | 0.0702 | 1.000 ± 0.1748 N=8 | 0.6031 ± 0.07778 N=7 |
| *SUMF2* | Cortex | 0.0321 | 1.000 ± 0.09093 N=8 | 0.6993 ± 0.08441 N=7 |
| *PMS2* | Cortex | 0.017 | 1.000 ± 0.1214 N=7 | 0.6348 ± 0.05137 N=7 |

N, sample size; SEM, standard error of the mean

**Supplementary Table 4** Differences in protein levels of top HD modifiers ascertained by Western blotting in HD and control brain microvascular endothelial cells (BMECs).

| Protein | *P*-value | Mean controls (SEM) | Mean HD (SEM) |
|---|---|---|---|
| CHCHD2 | 0.4491 | 1.000 ± 0.2146 N=6 | 0.7545 ± 0.2259 N=6 |
| EIF2AK1 | 0.0355 | 1.000 ± 0.1357 N=12 | 1.535 ± 0.1966 N=12 |
| FAN1 all | 0.6431 | 1.000 ± 0.06735 N=12 | 0.9434 ± 0.09993 N=12 |
| GPR161 | 0.526 | 1.000 ± 0.04335 N=11 | 1.070 ± 0.09644 N=12 |
| MSH3 | 0.0134 | 1.000 ± 0.06632 N=12 | 0.6667 ± 0.1089 N=10 |
| MTMR10 | 0.161 | 1.000 ± 0.05193 N=12 | 0.7575 ± 0.1589 N=12 |
| RRM2B | 0.0291 | 1.000 ± 0.03859 N=12 | 0.8124 ± 0.07049 N=12 |
| SUMF2 | 0.291 | 1.000 ± 0.08431 N=12 | 0.8720 ± 0.08294 N=12 |
| PMS2 | 0.2306 | 1.000 ± 0.1012 N=12 | 0.8151 ± 0.1101 N=11 |

N, sample size; SEM, standard error of the mean

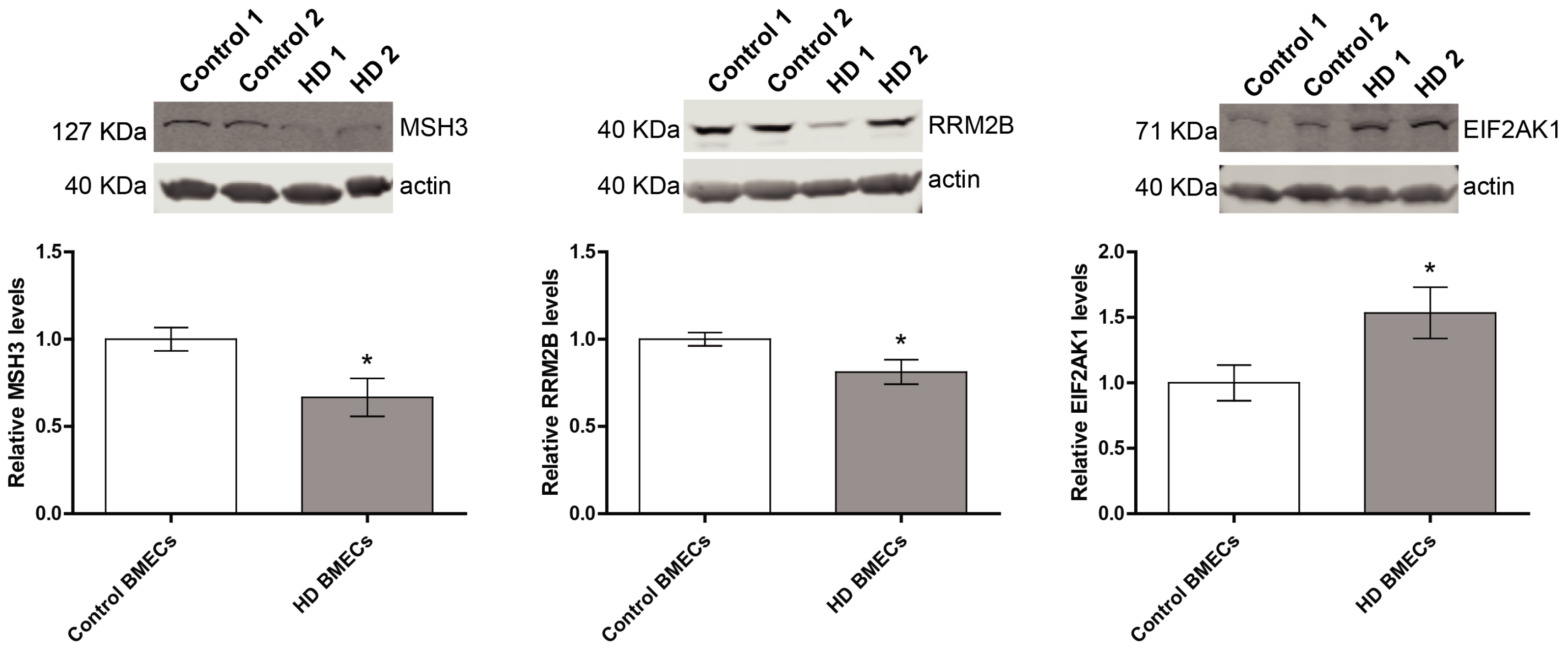


**Supplementary Fig. 2 Top candidate HD modifier proteins are dysregulated in brain microvascular endothelial cells (BMECs) derived from human pluripotent stems cells.** Relative protein levels of MSH3 and RRM2B are significantly decreased in HD patient derived BMECS compared to controls, while EIF2AK1 is significantly upregulated in HD BMECs. Asterisks indicate $P<0.05$.

**Supplementary Table 5** Enrichment analyses of S-PrediXcan associations for modifiers of the clinical onset of Huntington disease for diverse gene sets arranged by significance levels.

| Gene list | Description* | Reference(s)[a] | Top 5 genes[b] | Genes in set | Mean $Z^2$-score of gene set members | Mean $Z^2$-score of other genes | *P*-value | Corrected *P*-value |
|---|---|---|---|---|---|---|---|---|
| GWAS Catalog | Closest gene 3' and 5' of GWAS hits in the NHGRI GWAS Catalog as of Feb 9, 2015 | www.genome.gov/gwastudies PMID: 24316577 | *HCN4* *RBM6* *AKR1C4* *CYP27C1* *STX2* | 4081 | 0.95 | 0.97 | $7.6 \times 10^{-8}$ | **$1.3 \times 10^{-6}$** |
| Drug targets (Nelson *et al.*) | Drug targets according to Nelson *et al.* 2012 | PMID: 16376820 PMID: 22604722 | *CHRNA5* *NOS2* *GSK3B* *TRPM8* *CTLA4* | 201 | 1.18 | 0.97 | $1.7 \times 10^{-6}$ | **$2.9 \times 10^{-5}$** |
| DNA Repair Genes (Wood *et al.*) | Human DNA repair genes from Wood *et al.* | PMID: 15922366 | *MSH3* *PMS1* *PMS2* *ERCC3* *UBE2B* | 178 | 1.27 | 0.96 | $3.5 \times 10^{-5}$ | **0.00059** |
| ClinGen haplo-insufficient genes | Genes with sufficient evidence for dosage pathogenicity (level 3) as determined by the ClinGen Dosage Sensitivity Map | www.ncbi.nlm.nih.gov/projects/dbvar/clingen/ | *SETD5* *PTCH1* *TWIST1* *NR5A1* *PTPN11* | 221 | 1.08 | 0.97 | 0.00012 | **0.002** |
| Genes with any disease association reported in ClinVar | Genes where there is at least one variant with an assertion of pathogenic or likely pathogenic in ClinVar | PMID: 24234437 | *HCN4* *MSH3* *DPM2* *PMS1* *SQSTM1* | 3078 | 1.01 | 0.96 | 0.00041 | **0.007** |
| All autosomal dominant | Autosomal dominant genes according to | PMID: 22995991 | *HCN4* *FXYD2* *PMS1* | 709 | 1.09 | 0.96 | 0.0011 | **0.019** |

| | | | | | | | | |
|---|---|---|---|---|---|---|---|---|
| | either Berg *et al.* or Blekhman *et al.* | PMID: 18571414 | *SQSTM1* *PMS2* | | | | | |
| DNA Repair Genes (Kang *et al.*) | DNA repair genes from DNA repair pathways: ATM, BER, FA/HR, MMR, NHEJ, NER, TLS, XLR, RECQ, and other. | PMID: 22505474 | *MSH3* *PMS1* *PMS2* *ERCC3* *UBE2B* | 151 | 1.22 | 0.97 | 0.0032 | 0.054 |
| GPCRs | GPCR list from guidetopharmacology.org | PMID: 24234439 | *HCN4* *FXYD2* *KCNK16* *MST1R* *NR1I2* | 1705 | 0.94 | 0.97 | 0.0043 | 0.072 |
| Kinases | From UniProt (pkinfam) | PMID: 10647936 PMID: 12471243 PMID: 17557329 | *MST1R* *PHKG1* *CAMKV* *ULK3* *GRK6* | 351 | 0.94 | 0.97 | 0.0045 | 0.076 |
| All autosomal recessive | Autosomal recessive genes according to either Berg *et al.* or Blekhman *et al.* | PMID: 22995991 PMID: 18571414 | *ERCC3* *NDUFAF3* *AMT* *MPI* *ASH1L* | 1183 | 0.98 | 0.97 | 0.045 | 0.76 |
| Olfactory receptors | Olfactory receptors from the Mainland et al. 2015 | PMID: 25977809 | *OR52D1* *OR56B1* *OR6T1* *OR51S1* *OR51I2* | 371 | 0.87 | 0.97 | 0.047 | 0.8 |
| Natural product targets | List of hand-curated targets of natural products | PMID: 20565092 | *GSK3B* *MAP2K1* *JAK1* *PPIA* *POLD1* | 37 | 0.86 | 0.97 | 0.24 | 1 |

| | | | | | | | | |
|---|---|---|---|---|---|---|---|---|
| Essential in mice | Genes where homozygous knockout in mice results in pre-, peri- or post-natal lethality | PMID: 21051359 PMID: 23675308 PMID: 23843252 | *HCN4 NKX2-2 SRSF3 DDX20 ERCC3* | 2454 | 0.97 | 0.97 | 0.25 | 1 |
| BROCA Cancer Risk Panel | Panel used for the evaluation of patients with a suspected hereditary cancer predisposition | http://depts.washington.edu/labweb/Divisions/MolDiag/MolDiagGen/index.htm | *PMS2 PTCH1 CTNNA1 FLCN PDGFRA* | 66 | 1.25 | 0.97 | 0.36 | 1 |
| FDA-approved drug targets | Genes whose protein products are known to be the mechanistic targets of FDA-approved drugs | PMID: 24203711 PMID: 21059682 PMID: 18048412 PMID: 16381955 | *FXYD2 ATP4A TRPM8 P2RY12 CTLA4* | 286 | 0.96 | 0.97 | 0.4 | 1 |
| Essential in culture | Genes deemed essential in multiple cultured cell lines based on shRNA screen data | PMID: 24987113 | *SRSF3 NAPSA EIF2S2 HSPG2 NAPG* | 285 | 0.97 | 0.97 | 0.62 | 1 |
| ACMG V2.0 | Recommended for reporting of secondary findings in clinical sequencing by ACMG | PMID: 27854360 | *PMS2 BRCA2 TSC1 MUTYH BMPR1A* | 59 | 1.34 | 0.97 | 0.85 | 1 |

Significantly enriched gene lists after Bonferroni correction for multiple testing are indicated in bold. Abbreviations: ACMG, American College of Medical Genetics and Genomics; FDA, Food and Drug Administration; GPCR, G-protein-coupled receptor; GWAS, genome-wide association study; NHGRI, National Human Genome Research Institute; PMID, PubMed identifier. [a]Derived from https://github.com/macarthur-lab/gene_lists (downloaded November 2017) [b]According to absolute mean *Z*-score across all tissues.

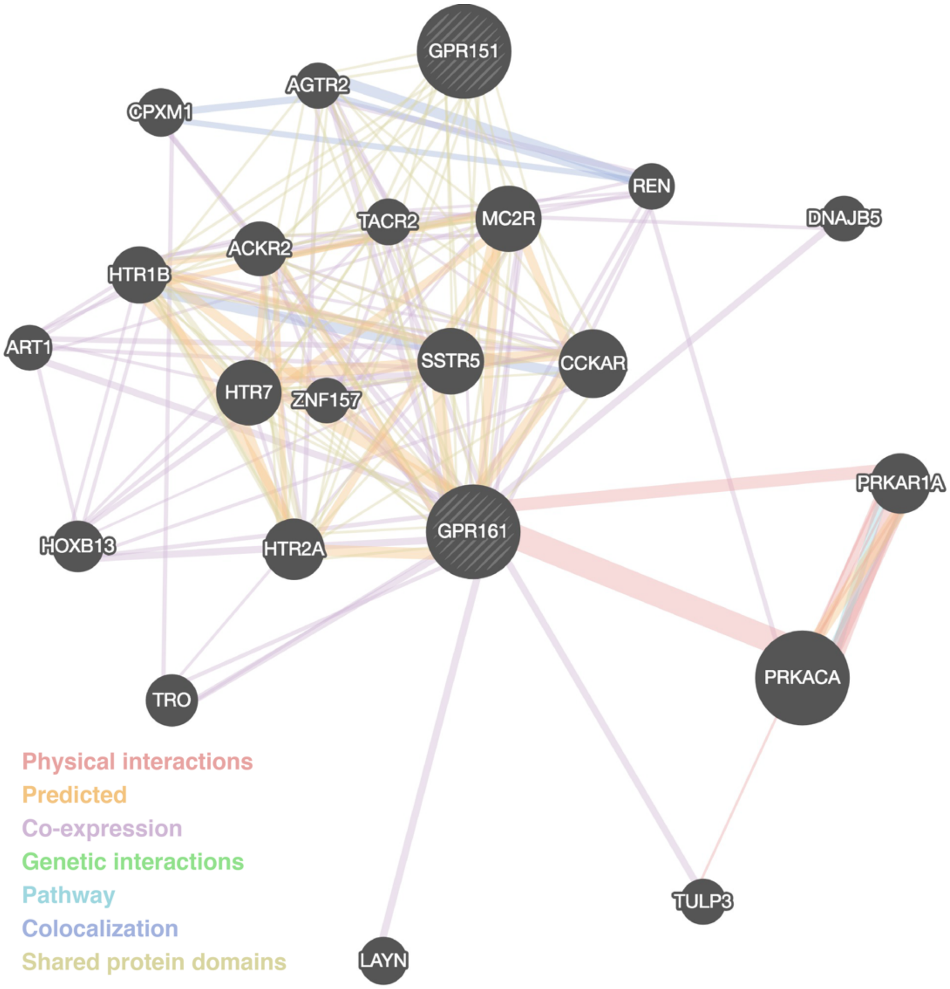


**Supplementary Fig. 3 Gene-gene functional interaction network for *GPR151* and *GPR161* to identify potential shared biological pathways between these two candidate HD clinical modifier genes.** GeneMania network analysis of these two G protein-coupled receptors indicated enrichment activation of phospholipase C activity and the serotonin receptor signaling pathway.

**Supplementary Table 6** Analysis of *Htt* CAG length and age dependent gene expression modules for enrichment of S-PrediXcan associations for modifiers of the clinical onset of Huntington disease. Striatal modules M1 and M20, both associated with increased expression at higher CAG lengths at increased ages were significantly enriched.

| Module | Number of genes in module | Top 5 genes[a] | Mean $Z^2$-score of module list members | Mean $Z^2$-score of other genes | *P*-value | Corrected *P*-value |
|---|---|---|---|---|---|---|
| Striatum M1 | 185 | *EIF2AK1* *TRAP1* *LRPPRC* *NAPG* *RNF14* | 1.26 | 0.96 | $8.70 \times 10^{-12}$ | **$1.60 \times 10^{-10}$** |
| Striatum M20 | 652 | *SUMF2* *MSH3* *FAM219B* *C22orf23* *ERCC3* | 1.08 | 0.96 | 0.00017 | **0.003** |
| Striatum M34 | 142 | *SRR* *ZNF667* *RUNX1T1* *ZSCAN20* *TBL1XR1* | 0.89 | 0.97 | 0.0065 | 0.12 |
| Cortex M4 | 282 | *ODF4* *RHOQ* *CINP* *GPR126* *CPNE9* | 0.92 | 0.97 | 0.0075 | 0.13 |
| Striatum M43 | 197 | *NICN1* *TEPP* *FUCA2* *POLR3C* *TTLL12* | 0.99 | 0.96 | 0.01 | 0.19 |
| Cortex M2 | 76 | *ANKRD63* *NFATC2* *EPYC* *SUSD2* | 1.03 | 0.97 | 0.039 | 0.7 |

| | | | | | | |
|---|---|---|---|---|---|---|
| | | *SH3RF2* | | | | |
| Striatum M52 | 79 | *CPT1A*<br>*OGDH*<br>*NT5C1A*<br>*FNIP2*<br>*PTPN22* | 0.87 | 0.97 | 0.2 | 1 |
| Cortex M6 | 226 | *HCN4*<br>*CPA6*<br>*FAM19A3*<br>*RD3L*<br>*SH3BGR* | 0.98 | 0.97 | 0.22 | 1 |
| Striatum M11 | 203 | *PHKG1*<br>*TENC1*<br>*DCN*<br>*JUN*<br>*PARP3* | 0.91 | 0.97 | 0.23 | 1 |
| Striatum M25 | 116 | *RHOQ*<br>*CD93*<br>*PRICKLE1*<br>*FLYWCH1*<br>*LRRFIP2* | 0.99 | 0.97 | 0.34 | 1 |
| Striatum M7 | 224 | *NKX2-2*<br>*LARS*<br>*HSPB6*<br>*SCCPDH*<br>*FGL2* | 0.96 | 0.97 | 0.43 | 1 |
| Striatum M2 | 1583 | *WRB*<br>*PARS2*<br>*AMT*<br>*ANKRD63* | 0.94 | 0.97 | 0.57 | 1 |
| Striatum M46 | 143 | *ZC3H14*<br>*ADNP*<br>*SLC35A3*<br>*FAM120B* | 0.92 | 0.97 | 0.66 | 1 |

| | | | | | | |
|---|---|---|---|---|---|---|
| | | *DHX57* | | | | |
| Striatum M39 | 245 | *AP3S2*<br>*DPYSL3*<br>*ANKRD61*<br>*ABCA7*<br>*PCDHB2* | 0.96 | 0.97 | 0.71 | 1 |
| Striatum M9 | 328 | *SQSTM1*<br>*ORMDL1*<br>*EIF2B2*<br>*ATP5G1*<br>*P4HTM* | 0.94 | 0.97 | 0.76 | 1 |
| Cortex M45 | 87 | *PHKG1*<br>*GRAMD3*<br>*C1orf110*<br>*DDAH1*<br>*CHPT1* | 0.99 | 0.97 | 0.77 | 1 |
| Striatum M10 | 145 | *CCT6A*<br>*GPM6A*<br>*PTPN9*<br>*KPNA4*<br>*KLHL7* | 0.90 | 0.97 | 0.81 | 1 |
| Cortex M7 | 101 | *CLIC4*<br>*GALNT6*<br>*PSAT1*<br>*MARCH8*<br>*TMEM88B* | 0.89 | 0.97 | 0.94 | 1 |

Significantly enriched modules after Bonferroni correction for multiple testing are highlighted in grey. [a]According to absolute mean *Z*-score across all tissues.

Gene functional enrichment analyses for significantly associated modules (from Langvelder et al. 2016):

**M1 striatum** IPA: Protein ubiquitination pathway; tRNA charging; Huntington's disease signaling.

Gene ontology (GO): Purine nucleotide binding; Cellular protein localization; Ubl conjugation pathway.

**M20 striatum** IPA: p53 signaling; *Brca1* in DNA damage response.

GO: Cell division; Protocadherin beta; Zinc finger, CH2H-like

=

**Supplementary Table 7** Analysis gene expression consensus modules (*n*=47) from postmortem dorsal lateral prefrontal cortex in ROSMAP participants (Mostafavi *et al.* 2018) identified 15 modules with significant differences S-PrediXcan mean $Z^2$-scores for modifiers of the clinical onset of Huntington disease. The table is sorted by strength of association with modules enriched in the Huntington disease related associations highlighted in grey.

| **Module** | **Number of genes in module** | **Top 5 genes**[a] | **Mean $Z^2$ score of module list members** | **Mean $Z^2$ score of other genes** | ***P*-value** | **Corrected *P*-value** |
|---|---|---|---|---|---|---|
| 107 | 376 | *RHOA*<br>*FAM219B*<br>*PFKFB2*<br>*TMEM100*<br>*SCRG1* | 0.84 | 0.97 | $1.1 \times 10^{-12}$ | $5.5 \times 10^{-11}$ |
| 109 | 335 | *NKX2-2*<br>*MED1*<br>*RHOQ*<br>*HEY2*<br>*FLCN* | 0.83 | 0.97 | $1.1 \times 10^{-11}$ | $5.2 \times 10^{-10}$ |
| 106 | 411 | *PHKG1*<br>*AMT*<br>*CCDC82*<br>*CMTM8*<br>*DBT* | 1.05 | 0.97 | $3.1 \times 10^{-9}$ | $1.5 \times 10^{-7}$ |
| 108 | 255 | *TRIOBP*<br>*PTCH1*<br>*TUFM*<br>*TMEM216*<br>*NWD1* | 1.11 | 0.97 | $2.5 \times 10^{-7}$ | $1.2 \times 10^{-5}$ |
| 1 | 427 | *DPM2*<br>*P4HTM*<br>*LSMD1*<br>*LSM7*<br>*SH3GLB2* | 0.89 | 0.97 | $3.6 \times 10^{-7}$ | $1.8 \times 10^{-5}$ |

| | | | | | | |
|---|---|---|---|---|---|---|
| 114 | 232 | *STX2*<br>*DLL4*<br>*MYL6*<br>*SH3BGR*<br>*HEATR2* | 1.11 | 0.97 | $2.1 \times 10^{-6}$ | 0.0001 |
| 131 | 25 | *MDH1*<br>*SEC31A*<br>*TOMM70A*<br>*CD200*<br>*CMAS* | 1.43 | 0.97 | $4.1 \times 10^{-6}$ | 0.0002 |
| 257 | 37 | *CYB561D2*<br>*IFT52*<br>*DOPEY1*<br>*PIGT*<br>*POLDIP2* | 0.79 | 0.97 | $1.5 \times 10^{-5}$ | 0.0007 |
| 14 | 277 | *ATP5G1*<br>*DSCAML1*<br>*ANKRD39*<br>*ASNS*<br>*BNIP1* | 0.85 | 0.97 | $3.7 \times 10^{-5}$ | 0.002 |
| 233 | 39 | *MRPS18A*<br>*EIF1AD*<br>*BRK1*<br>*HMGA1*<br>*ZNF622* | 0.68 | 0.97 | $4.1 \times 10^{-5}$ | 0.002 |
| 16 | 305 | *CAMKV*<br>*C1QTNF4*<br>*ZER1*<br>*FBLL1*<br>*FAM212B* | 1 | 0.97 | $4.5 \times 10^{-5}$ | 0.002 |
| 122 | 306 | *EIF2AK1*<br>*PMS2*<br>*ERCC3*<br>*QRICH1*<br>*CCZ1* | 1.16 | 0.96 | $4.8 \times 10^{-5}$ | 0.002 |

| | | | | | | |
|---|---|---|---|---|---|---|
| 117 | 373 | *SRSF3*<br>*ASNSD1*<br>*EIF2B2*<br>*TIGD7*<br>*MOB1B* | 0.89 | 0.97 | $1.9 \times 10^{-4}$ | 0.009 |
| 11 | 13 | *DCLRE1C*<br>*CNOT2*<br>*ZNF232*<br>*RHOU*<br>*ADAL* | 0.66 | 0.97 | $2.0 \times 10^{-4}$ | 0.010 |
| 2 | 412 | *SUMF2*<br>*ZNF75A*<br>*AP3S2*<br>*MAP3K13*<br>*CCZ1B* | 1.02 | 0.97 | 0.0003 | 0.013 |

Significantly enriched modules after Bonferroni correction for multiple testing are highlighted in grey.
[a]According to absolute mean *Z*-score across all tissues.

**Supplementary Table 8** Associations between ROSMAP cortical coexpression modules and phenotypes that passed within trait Bonferroni correction for multiple testing.

| Module | Trait | Category | *N* | Estimate | SE | Statistic | *P*-value |
|---|---|---|---|---|---|---|---|
| m131 | Estimated slope from random effects model for working memory | Cognitive Decline | 470 | 0.35 | 0.06 | 5.91 | 6.72E-09 |
| m131 | Mini-Mental State Exam, 30 item | Cognitive Decline | 494 | 53 | 10 | 5.12 | 4.46E-07 |
| m122 | Global burden of AD pathology based on 5 regions | Pathology | 494 | -5.8 | 1.2 | -4.9 | 1.31E-06 |
| m131 | Global cognitive function - Average of 19 tests | Cognitive Decline | 493 | 6.5 | 1.3 | 4.89 | 1.35E-06 |
| m131 | Estimated slope from random effects model for perceptual speed | Cognitive Decline | 457 | 0.43 | 0.094 | 4.57 | 6.41E-06 |
| m114 | Estimated slope from random effects model for working memory | Cognitive Decline | 470 | -0.37 | 0.086 | -4.31 | 2.02E-05 |
| m122 | Estimated slope from random effects model for semantic memory | Cognitive Decline | 461 | 1 | 0.23 | 4.3 | 2.08E-05 |
| m114 | Estimated slope from random effects model for perceptual speed | Cognitive Decline | 457 | -0.57 | 0.13 | -4.3 | 2.13E-05 |
| m114 | Mini-Mental State Exam, 30 item | Cognitive Decline | 494 | -63 | 15 | -4.26 | 2.47E-05 |
| m131 | Estimated slope from random effects model for global cognition | Cognitive Decline | 470 | 0.52 | 0.12 | 4.25 | 2.60E-05 |
| m122 | NIA Reagan Diagnosis of Alzheimer's disease - 4 levels (none to high likelihood) | Pathology | 494 | 6 | 1.4 | 4.23 | 2.78E-05 |
| m122 | Global cognitive function - Average of 19 tests | Cognitive Decline | 493 | 9.1 | 2.2 | 4.18 | 3.48E-05 |
| m122 | Estimated slope from random effects model for global cognition | Cognitive Decline | 470 | 0.84 | 0.2 | 4.18 | 3.50E-05 |
| m122 | Neuritic plaque summary based on 5 regions | Pathology | 494 | -26 | 6.2 | -4.14 | 4.07E-05 |
| m122 | Diffuse plaque summary based on 5 regions | Pathology | 494 | -26 | 6.5 | -4.1 | 4.92E-05 |

| | | | | | | | |
|---|---|---|---|---|---|---|---|
| m108 | Estimated slope from random effects model for working memory | Cognitive Decline | 470 | -0.32 | 0.078 | -4.08 | 5.40E-05 |
| m122 | Estimated slope from random effects model for perceptual speed | Cognitive Decline | 457 | 0.62 | 0.15 | 4 | 7.38E-05 |
| m131 | Parkinsonian signs domain: Gait | Motor and Gait | 491 | -100 | 26 | -3.88 | 0.00012 |
| m16 | Motor function partial composite: Hand strength | Motor and Gait | 465 | 1.7 | 0.45 | 3.81 | 0.00016 |
| m114 | Global cognitive function - Average of 19 tests | Cognitive Decline | 493 | -7.1 | 1.9 | -3.73 | 0.000215 |
| m122 | Mini-Mental State Exam, 30 item | Cognitive Decline | 494 | 63 | 17 | 3.72 | 0.000223 |
| m122 | Estimated slope from random effects model for episodic memory | Cognitive Decline | 469 | 0.76 | 0.21 | 3.7 | 0.00024 |
| m114 | Estimated slope from random effects model for global cognition | Cognitive Decline | 470 | -0.64 | 0.17 | -3.67 | 0.00027 |
| m106 | Motor function partial composite: Hand strength | Motor and Gait | 465 | -2.1 | 0.57 | -3.64 | 0.000304 |
| m16 | Perceived social isolation (loneliness) | Lifestyle, Personality | 202 | -5 | 1.4 | -3.58 | 0.000428 |
| m108 | Motor function partial composite: Hand strength | Motor and Gait | 465 | -1.7 | 0.48 | -3.44 | 0.00064 |
| m131 | Estimated slope from random effects model for episodic memory | Cognitive Decline | 469 | 0.43 | 0.13 | 3.43 | 0.000667 |
| m131 | Perceived social isolation (loneliness) | Lifestyle, Personality | 202 | -4.3 | 1.2 | -3.45 | 0.00069 |
| m131 | Global burden of AD pathology based on 5 regions | Pathology | 494 | -2.4 | 0.74 | -3.31 | 0.00101 |
| m114 | Perceived social isolation (loneliness) | Lifestyle, Personality | 202 | 6.1 | 1.8 | 3.34 | 0.00102 |
| m131 | Estimated slope from random effects model for semantic memory | Cognitive Decline | 461 | 0.47 | 0.14 | 3.25 | 0.00122 |
| m108 | Mini-Mental State Exam, 30 item | Cognitive Decline | 494 | -44 | 13 | -3.25 | 0.00123 |
| m114 | NIA Reagan Diagnosis of Alzheimer's disease - 4 levels (none to high likelihood) | Pathology | 494 | -4 | 1.2 | -3.22 | 0.00138 |

| | | | | | | | |
|---|---|---|---|---|---|---|---|
| m114 | Estimated slope from random effects model for semantic memory | Cognitive Decline | 461 | -0.65 | 0.2 | -3.18 | 0.00157 |
| m114 | Estimated slope from random effects model for episodic memory | Cognitive Decline | 469 | -0.57 | 0.18 | -3.17 | 0.00162 |
| m131 | NIA Reagan Diagnosis of Alzheimer's disease - 4 levels (none to high likelihood) | Pathology | 494 | 2.7 | 0.87 | 3.14 | 0.00177 |
| m108 | Global cognitive function - Average of 19 tests | Cognitive Decline | 493 | -5.4 | 1.7 | -3.14 | 0.00182 |
| m114 | Actical sleep measure - Probability of transitioning from rest state to active state | Sleep and Circadian Rhythms | 95 | 0.13 | 0.04 | 3.17 | 0.00204 |
| m131 | Neuritic plaque summary based on 5 regions | Pathology | 494 | -12 | 3.8 | -3.05 | 0.00242 |
| m131 | Diffuse plaque summary based on 5 regions | Pathology | 494 | -12 | 4 | -3.02 | 0.00262 |
| m122 | Estimated slope from random effects model for working memory | Cognitive Decline | 470 | 0.3 | 0.1 | 2.99 | 0.00291 |
| m114 | Neuritic plaque summary based on 5 regions | Pathology | 494 | 16 | 5.4 | 2.98 | 0.00303 |
| m114 | Diffuse plaque summary based on 5 regions | Pathology | 494 | 17 | 5.7 | 2.96 | 0.00321 |
| m114 | Global burden of AD pathology based on 5 regions | Pathology | 494 | 3 | 1.1 | 2.86 | 0.00441 |

**Supplementary Table 9** Associations between expression of top transcriptomic modifier genes in the ROSMAP cortical samples and phenotypes that passed within trait Bonferroni correction for multiple testing.

| **Gene** | **Trait** | **Signed -log$_{10}$(*P*-value)** | ***P*-value** |
|---|---|---|---|
| *MTMR10* | Global cognitive function decline | -6.91 | 1.24E-07 |
| *PMS2* | Global cognitive function decline | 5.89 | 1.28E-06 |
| *MTMR10* | Episodic memory decline | -5.87 | 1.35E-06 |
| *MTMR10* | Clinical AD diagnosis 2 | 5.83 | 1.48E-06 |
| *MTMR10* | Amyloid plaque | 5.41 | 3.86E-06 |
| *MTMR10* | Clinical diagnosis | 5.27 | 5.32E-06 |
| *MTMR10* | Post mortem clinical AD diagnosis | 4.82 | 1.50E-05 |
| *PMS2* | Episodic memory decline | 4.58 | 2.63E-05 |
| *MTMR10* | Neurotic plaque | 3.47 | 0.000341 |
| *PMS2* | Neurotic plaque | -3.40 | 0.000398 |
| *PMS2* | Amyloid plaque | -3.17 | 0.000684 |
| *CHCHD2* | Global cognitive function decline | 3.15 | 0.000708 |
| *CHCHD2* | Neurotic plaque | -2.76 | 0.00172 |
| *PMS2* | Clinical AD diagnosis 2 | -2.73 | 0.00188 |
| *PMS2* | Clinical diagnosis | -2.53 | 0.00292 |
| *CHCHD2* | Episodic memory decline | 2.43 | 0.0036 |

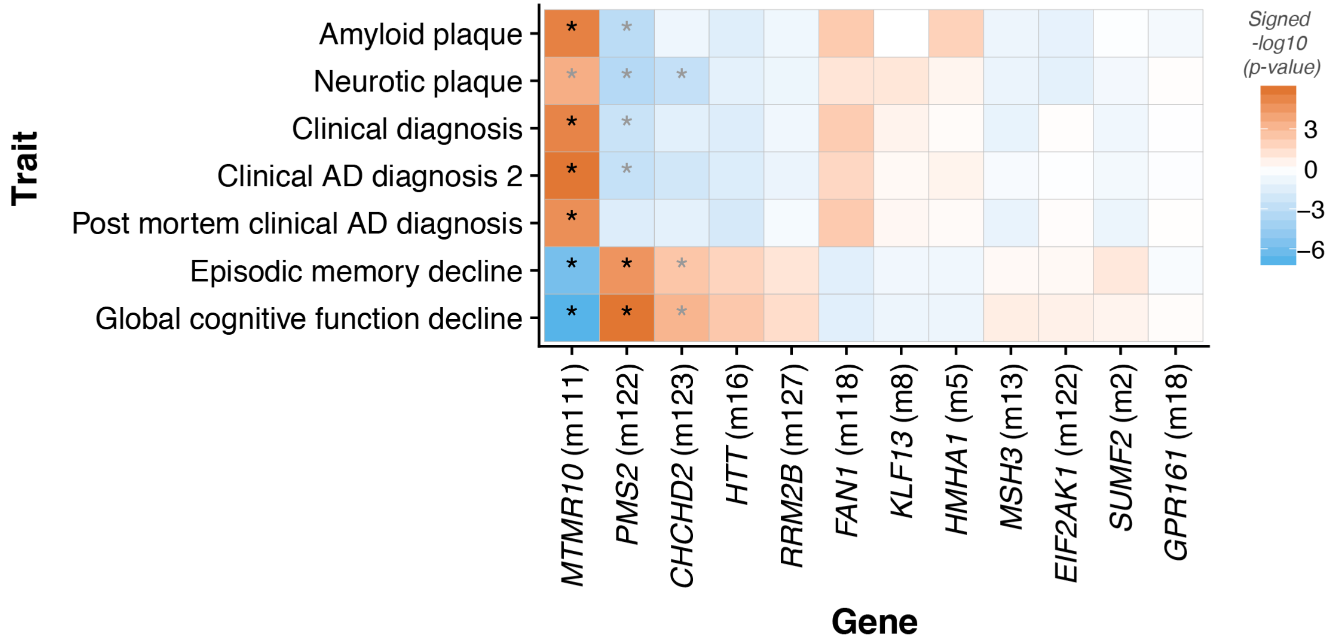


**Supplementary Fig. 4** Analysis of top transcriptomic modifier genes for Huntington disease clinical onset, as well as *HTT*, with regards to a subset of the phenotypes. Expression of *MTMR10* and *PMS2* was associated with a variety of phenotypes, decline in global cognitive function achieving greatest statistical significance. Black asterisks: significant after correcting for all comparisons, grey asterisks: significant after adjusting for within trait comparisons. AD, Alzheimer's disease.

**Supplementary Table 10. Top perturbagens similar (>95%) and dissimilar (<95%) perturbagens in the signature matching analyses of HD gene expression TWAS profiles to the Connectivity Map database**

| Type | Name | Median tau | PC3 | VCAP | A375 | A549 | HA1E | HCC515 | HT29 | MCF7 | HEPG2 | Summary |
|---|---|---|---|---|---|---|---|---|---|---|---|---|
| CP | CGP-60474 | 99.69 | 99.56 | 99.96 | 99.63 | 99.74 | 98.42 | 99.73 | 99.85 | 99.65 | 99.74 | 99.05 |
| KD | SLC25A28 | 99.67 | 99.82 | NA | 86.75 | 67.58 | 99.73 | NA | 65.72 | 99.61 | 99.95 | 99.92 |
| CP | chromomycin-a3 | 99.64 | 99.71 | 60.31 | 99.81 | 99.19 | 99.65 | 99.54 | 99.63 | 99.76 | 99.86 | 99.01 |
| CP | ER-27319 | 99.62 | 99.66 | 99.34 | 91.77 | 99.63 | 99.61 | 99.62 | 99.61 | 99.62 | 99.63 | 98.73 |
| CP | mitoxantrone | 99.6 | 99.6 | 99.03 | 99.63 | 83.06 | 99.6 | 99.75 | 99.76 | 99.43 | NA | 98.87 |
| CP | alvocidib | 99.59 | 99.56 | 92.28 | 99.66 | 99.64 | 99.64 | 99.34 | 99.63 | 99.48 | 99.66 | 98.66 |
| CP | daunorubicin | 99.58 | 99.24 | 99.85 | 99.65 | 99.76 | 98.85 | 99.87 | 99.6 | 99.53 | 99.56 | 98.63 |
| CP | daunorubicin | 99.57 | 99.6 | 95.74 | 99.65 | 99.79 | 99.58 | 99.45 | 99.6 | 99.55 | 99.42 | 98.63 |
| CP | BRD-K77681376 | 99.56 | 99.34 | NA | 99.58 | 99.66 | NA | 99.53 | 99.65 | 98.14 | 99.66 | 98.66 |
| CP | bisindolylmaleimide-ix | 99.56 | 99.22 | 69.45 | 99.72 | 99.55 | 99.57 | 99.57 | 99.68 | 99.49 | 99.56 | 98.63 |
| CP | triptolide | 99.56 | 99.63 | 99.93 | 97.25 | 99.68 | 99.51 | 99.61 | 99.44 | 99.48 | 99.67 | 98.59 |
| CP | AT-7519 | 99.54 | 99.52 | 98.8 | 99.67 | 99.56 | 99.68 | 99.63 | 99.9 | 99.42 | 99.4 | 98.77 |
| CP | PIK-75 | 99.53 | 99.6 | -91.14 | 99.81 | 97.72 | 99.87 | 99.51 | 99.61 | 99.53 | 99.54 | 98.66 |
| CP | JNK-9L | 99.53 | 99.66 | 97.16 | 99.47 | 99.62 | 99.68 | 99.51 | 99.55 | 99.26 | 99.7 | 98.66 |
| CP | doxorubicin | 99.52 | 99.52 | 85.76 | 99.95 | 93.95 | 99.65 | 99.63 | 86.24 | 99.72 | NA | 98.94 |
| CP | daunorubicin | 99.52 | 99.57 | 99.19 | 99.72 | 94.53 | 99.78 | 99.54 | 99.5 | 99.27 | 99.73 | 98.8 |
| CP | pidorubicine | 99.52 | 99.65 | 84.87 | 99.5 | 85.74 | 99.64 | 99.5 | 99.55 | 99.81 | 99.54 | 98.7 |
| CP | triptolide | 99.51 | 99.52 | 60.34 | 99.5 | 99.2 | 99.68 | NA | 99.93 | 99.68 | NA | 98.87 |
| CP | dactinomycin | 99.51 | 99.53 | 89.19 | 99.88 | 95.14 | 99.51 | 99.56 | 99.51 | 99.59 | 99.37 | 98.63 |
| CP | PF-562271 | 99.51 | 99.63 | NA | 99.65 | 99.25 | 99.61 | 99.51 | 99.5 | 95.99 | 99.54 | 98.63 |
| CP | ZG-10 | 99.51 | 99.53 | 99.87 | 99.59 | 95.02 | 99.64 | 99.48 | 99.64 | 99.49 | 99.46 | 98.56 |
| CP | A-443644 | 99.43 | 99.66 | 61.92 | 95.42 | 99.73 | 99.38 | 99.69 | 99.59 | 72.21 | 99.48 | 98.66 |
| CP | dactinomycin | 99.43 | 99.62 | 92.65 | 99.39 | 99.62 | 99.77 | 97.84 | 99.73 | 93.78 | 99.47 | 98.56 |
| CP | BRD-K85853281 | 99.4 | 99.59 | 80.7 | 99.42 | 99.66 | 91.83 | 99.37 | 99.68 | 99.32 | 99.54 | 98.63 |
| KD | PLS1 | 99.38 | 99.92 | NA | 99.98 | 98.61 | -86.05 | 99.96 | 30.51 | 99.38 | 90.58 | 99.95 |

| | | | | | | | | | | | | |
|---|---|---|---|---|---|---|---|---|---|---|---|---|
| KD | NFKBIB | 99.37 | 99.98 | 100 | 99.62 | 94.05 | 96.27 | 86.29 | 99.61 | 99.13 | 99.12 | 99.82 |
| CP | NVP-BEZ235 | 99.22 | 99.1 | 99.34 | 99.56 | 88.61 | 99.46 | 99.44 | 99.81 | 85.99 | 32.58 | 98.84 |
| CP | lestaurtinib | 99.19 | 99.66 | 99.91 | 85.7 | 99.86 | 98.89 | 99.5 | 98.81 | 97.43 | 99.5 | 98.7 |
| CP | JNJ-7706621 | 99.18 | 99.45 | 97.25 | 99.59 | 77.57 | 93.51 | 98.92 | 99.56 | 99.53 | 99.59 | 98.52 |
| CP | BRD-K53780220 | 99.17 | 92.01 | 88.67 | 99.17 | 32.89 | NA | 99.54 | 99.79 | 99.78 | 99.77 | 98.84 |
| CP | idarubicin | 99.16 | 99.5 | 90.32 | 99.67 | 99.33 | 99.58 | NA | 98.99 | 89.65 | NA | 98.63 |
| CP | AZD-8055 | 99.08 | 99.8 | 99.43 | 99.89 | 74.58 | 99.11 | 99.57 | 98.52 | 99.05 | 93.96 | 98.79 |
| CP | KU-0063794 | 99.03 | 99.11 | 99.67 | 97.07 | 50.14 | 98.42 | 99.69 | 99.25 | 99.76 | -56.51 | 98.94 |
| PCL | PI3K inhibitor | 99 | 99.17 | 99.45 | 96.47 | 90.47 | 99.5 | 99.2 | 98.83 | 96.42 | 85.11 | 99.2 |
| PCL | Topoisomerase inhibitor | 98.98 | 99.49 | 89.67 | 98.62 | 91.59 | 96.78 | 99.57 | 99.4 | 95.6 | 99.52 | 99.33 |
| CP | BMS-345541 | 98.92 | 43.56 | 95.19 | -59.08 | 99.79 | 99.63 | 99.44 | 99.08 | -93.03 | 99.75 | 98.77 |
| KD | NUCKS1 | 98.91 | 94.97 | 45.69 | 99.66 | 65.91 | 83.06 | NA | 98.91 | 99.7 | 99.66 | 99.66 |
| PCL | MTOR inhibitor | 98.9 | 98.99 | 99.39 | 95.1 | 83.65 | 99.29 | 99.5 | 98.82 | 98.73 | -72.56 | 99.22 |
| CP | pirarubicin | 98.87 | 98.87 | 78.79 | 99.49 | 93.09 | 96.55 | 99.47 | 98.87 | 99.71 | 99.44 | 98.38 |
| CP | AS-601245 | 98.85 | 99.52 | 96.96 | 99.72 | 92.27 | 92.1 | 99.26 | 99.48 | 78.81 | 99.43 | 98.45 |
| KD | PLAT | 98.82 | 99.01 | 99.21 | 35.64 | 67.89 | 56.61 | NA | 37.43 | 98.91 | 98.82 | 99 |
| KD | PLAUR | 98.66 | -60 | NA | 41.68 | NA | 98.85 | NA | 98.66 | 99.9 | 64.67 | 98.87 |
| CP | 5-iodotubercidin | 98.65 | 99.89 | 81.51 | 99.8 | 98.73 | 88.77 | 99.41 | 99.62 | 76.17 | 94.22 | 98.56 |
| KD | CTDSP1 | 98.64 | -36 | 69.44 | 99.11 | 99.95 | 98.64 | -71.1 | -20.77 | NA | 99.81 | 99.82 |
| CP | ISOX | 98.52 | 85.03 | 98.84 | 94.62 | 98.24 | 94.99 | 99.25 | 99.14 | 50.64 | 99.5 | 98.8 |
| KD | IL10RB | 98.36 | 84.15 | NA | 66.49 | NA | 99.87 | NA | 98.36 | 99.75 | 65.38 | 99.89 |
| KD | TNIP1 | 98.32 | -28.03 | 54.95 | -29.73 | 83.22 | 97.63 | 99.71 | 99.82 | 99.02 | 99.58 | 99.71 |
| CP | PI-828 | 98.29 | 99.72 | 99.5 | 98.38 | 89.66 | 98.47 | 99.85 | 90.04 | 70.04 | 43.13 | 98.2 |
| CP | apicidin | 98.24 | 77.62 | 52.07 | 93.34 | 98.99 | 99.31 | 98.92 | 99.38 | 32.02 | 98.6 | 97.89 |
| PCL | CDK inhibitor | 98.2 | 99.95 | 66.89 | 92.12 | 73.37 | 99.54 | 99.44 | 97.24 | 99.17 | 84.96 | 99.55 |
| CP | PHA-793887 | 98.15 | 99.65 | 58.51 | 95.71 | 69.02 | 98.81 | 98.19 | 99.5 | 99.09 | 84.92 | 98.1 |
| CP | staurosporine | 98.06 | 99.58 | 81.28 | 94.72 | 99.63 | 97.55 | 99.47 | 99.56 | 97.01 | 97.05 | 98.56 |
| CP | purvalanol-a | 97.99 | 98.4 | 90.69 | 24.29 | 66.23 | 98.98 | 99.18 | -38.76 | 99.71 | NA | 97.99 |
| CP | doxorubicin | 97.97 | 99.57 | 77.44 | 98.62 | 69.48 | 99.52 | 99.9 | 97.59 | 91.56 | 97.64 | 98.31 |

| | | | | | | | | | | | | |
|---|---|---|---|---|---|---|---|---|---|---|---|---|
| KD | GTF2A2 | 97.82 | 99.53 | -90.1 | -72.83 | 99.15 | 99.73 | NA | 42.71 | -18.11 | 97.82 | 99.82 |
| KD | GMPS | 97.81 | 97.45 | 98.16 | 99.26 | 83.02 | 91.75 | 96.39 | 98.94 | 99.79 | 97.13 | 99.55 |
| CP | TPCA-1 | 97.72 | 96.2 | 99.91 | 98.42 | 45.83 | 98.42 | 98.57 | 97.67 | -49.3 | 40.3 | 97.78 |
| CP | AZD-7762 | 97.69 | 99.35 | 77.78 | 79.5 | 99.35 | 98.95 | 78.12 | 97.3 | 98.07 | 77.1 | 98.27 |
| KD | PSEN1 | 97.67 | 99.96 | 97.67 | 79.38 | 99.93 | 44.07 | 62.69 | 99.96 | NA | 20.26 | 99.95 |
| CP | fludarabine | 97.67 | 99.12 | NA | 96.93 | 90.59 | 95.51 | 98.72 | 98.93 | 93.04 | NA | 98.41 |
| KD | RALBP1 | 97.42 | 99.35 | 97.26 | 63.47 | 99.13 | 97.58 | 49.89 | 93.18 | 99.81 | 81.93 | 98.98 |
| CP | L-690488 | 97.26 | 96.43 | 96.78 | 92.19 | 37.92 | 99.31 | 97.73 | 99.94 | 20.01 | 99.66 | 99.08 |
| KD | RPS3 | 97.17 | 97.38 | -99.14 | 96.97 | 99.78 | 84.9 | 65.08 | 99.23 | -35.46 | 99.96 | 99.31 |
| CP | fostamatinib | 97.15 | 99.77 | 96.06 | 98.23 | 99.09 | 91.07 | 95.95 | 99.91 | -85.51 | -66.63 | 98.98 |
| KD | RASA1 | 97.11 | 39.86 | NA | 98.13 | 31.33 | 37.65 | NA | NA | 98.32 | 97.11 | 99.53 |
| KD | ZNF274 | 97.05 | 86.5 | 23.79 | 97.05 | -83.49 | 56.82 | 99.34 | NA | 99.83 | 97.85 | 99.61 |
| CP | PI-103 | 97.03 | 98.65 | 97.29 | 93.43 | 96.78 | 99.19 | 99.11 | 44.72 | 86.73 | 82.72 | 98.29 |
| KD | IL2RB | 96.89 | 99.35 | 100 | 63.14 | 68.2 | 99.82 | 50.66 | 70.18 | 96.89 | NA | 99.61 |
| KD | UVRAG | 96.84 | -94.38 | NA | 96.84 | 99.29 | 99.98 | -35.08 | 30 | 99.91 | 34.38 | 99.79 |
| PCL | FLT3 inhibitor | 96.81 | 99.69 | 97.86 | 91.36 | 91.88 | 96.86 | 99.81 | 94.08 | 99.37 | -30.1 | 96.76 |
| KD | NPTN | 96.73 | 37.76 | 52.8 | 96.73 | 87.82 | 99.67 | 99.4 | 99.32 | 85.84 | NA | 99.53 |
| KD | CBX6 | 96.69 | 96.69 | 81.2 | 22.24 | 96.43 | 99.89 | NA | 99.6 | 92.44 | 99.32 | 99.87 |
| CP | U-0126 | 96.69 | 74.39 | 69.85 | 99.55 | 97.02 | 59.17 | 87.44 | 96.37 | 99.65 | 97.84 | 99.21 |
| CP | temsirolimus | 96.58 | 99.33 | 98.21 | 93.33 | 96.11 | 99 | 98.82 | 76.76 | 91.7 | -40.57 | 97.04 |
| KD | MED28 | 96.5 | 96.99 | NA | 95.84 | 27.88 | 96.01 | NA | 98.95 | 99.9 | -39.55 | 98.82 |
| CP | topotecan | 96.48 | 99.45 | 95.81 | 98.25 | 70.99 | 36.68 | 99.48 | 96.48 | 90.29 | NA | 97.57 |
| KD | CREBL2 | 96.47 | 92.23 | NA | 96.85 | 99.95 | 79.56 | NA | 91.06 | 99.36 | 96.08 | 98.67 |
| KD | TARDBP | 96.44 | 93.8 | 94.05 | 71.03 | 99.55 | 99.86 | 98.84 | 99.98 | 84.23 | 60.57 | 99.92 |
| CP | trichostatin-a | 96.44 | 91.81 | 81.8 | 96.63 | 98.77 | 98.23 | 96.25 | 91.7 | 75.96 | 99.13 | 97.08 |
| KD | ZNF219 | 96.35 | 51.3 | -95.84 | 93.44 | 98.59 | 97.56 | NA | 96.83 | 78.79 | 98.23 | 96.35 |
| KD | NR2C2 | 96.02 | 96.02 | NA | 53.58 | 38.69 | 97.97 | NA | NA | 98.43 | 87.34 | 97.29 |
| KD | ETV1 | 96.01 | 28.55 | 48.08 | 38.16 | 99.76 | 99.34 | 33.01 | 99.45 | 93.28 | 98.73 | 99.84 |
| CP | doxorubicin | 96 | 93.86 | 81.04 | 95.74 | 96.26 | 76.99 | 92.94 | 98.78 | 98.12 | 98.39 | 97.18 |

| | | | | | | | | | | | | |
|---|---|---|---|---|---|---|---|---|---|---|---|---|
| KD | CLCN5 | 95.98 | 98.02 | 100 | 93.82 | 93.94 | 80.13 | NA | -23.74 | NA | 99.25 | 99.45 |
| KD | ERGIC2 | 95.96 | 99.54 | 99.8 | -33.76 | 92.62 | 28.14 | 85.78 | NA | 99.51 | NA | 99.29 |
| KD | P4HA1 | 95.84 | 97.24 | 95.39 | 99.4 | 91.02 | 62.93 | 99.04 | 78.64 | 95.84 | NA | 99.53 |
| KD | EIF2AK2 | 95.7 | 97.41 | NA | 98.62 | NA | -93.74 | NA | 29.45 | 95.7 | 86.95 | 98.11 |
| OE | PRAME | 95.68 | 52.2 | 99.3 | -26.5 | 92.73 | 99.96 | 98.63 | 99.96 | -82.67 | -81.88 | 99.91 |
| KD | TACC3 | 95.68 | 63.25 | 27.85 | 99.36 | 97.86 | 99.12 | NA | 26 | 95.68 | 48.53 | 99.28 |
| CP | PP-30 | 95.68 | 93.71 | 99.91 | 97.65 | 71.92 | 75.55 | 84.88 | 99.66 | 99.42 | 67.83 | 98.73 |
| KD | ATP1A3 | 95.68 | 95.68 | 97.65 | 58.72 | 61.95 | 74.28 | NA | 97.75 | 91.58 | 97.89 | 97.8 |
| KD | FZD5 | 95.61 | 98.78 | 100 | 84.3 | -65.07 | 95.73 | 95.48 | 99.98 | 77.79 | -76.59 | 98.97 |
| KD | IRAK1 | 95.55 | 95.8 | -99.17 | 74.69 | NA | 99.2 | NA | 20.76 | 95.29 | 97.9 | 97.61 |
| CP | methotrexate | 95.42 | 97.47 | 99.26 | 37.08 | 82.81 | -45.53 | NA | 97.75 | 93.37 | NA | 97.78 |
| CP | guanabenz | 95.41 | 36.56 | 98.62 | -32.95 | -83.85 | 91.62 | 95.55 | 99.04 | 95.41 | NA | 98.13 |
| KD | PSEN2 | 95.26 | -41.11 | 34.83 | NA | 94.1 | 98.26 | 99.06 | 26.17 | 96.43 | NA | 98.47 |
| KD | ZNF791 | -95.11 | -91.8 | -46.84 | -99.91 | 39.96 | -98.6 | -93.15 | -99.45 | -97.07 | -36.98 | -99.71 |
| KD | LEPRE1 | -95.21 | -99.22 | 83.02 | NA | -99.3 | -67.41 | NA | -77.69 | -96.48 | -93.94 | -99.5 |
| KD | UGT1A6 | -95.26 | 62.74 | -95.26 | NA | -98.59 | 99.82 | -97.88 | -50.79 | -34.29 | -99.56 | -97.74 |
| KD | IRS2 | -95.34 | -58.82 | NA | -20.09 | -95.34 | -94.75 | NA | NA | -97.87 | -97.13 | -97.79 |
| KD | UBE2C | -95.38 | -98.95 | -38.59 | -93.62 | -94.71 | -91.75 | -96.04 | -99.13 | -98.77 | -20.85 | -98.44 |
| KD | SLC7A11 | -95.56 | -91.74 | -99.39 | -91.85 | -84.24 | -99.48 | NA | NA | NA | NA | -99.28 |
| KD | NCF2 | -95.67 | -99.17 | NA | 55.7 | -82.34 | -99.19 | NA | -96.57 | -94.78 | -41.77 | -99.76 |
| KD | GPATCH8 | -95.69 | NA | NA | -97.25 | -54 | -99.58 | -94.13 | -99.16 | -61.54 | -84.48 | -99.74 |
| KD | GCDH | -95.8 | -99.98 | 41.54 | 24.69 | -95.8 | -96.45 | -98.47 | -60.53 | NA | -92.78 | -98.47 |
| KD | FOXA1 | -95.85 | -99.52 | -95.85 | 44.77 | -96.81 | -68.75 | -99.56 | -51.33 | -25.38 | NA | -99.35 |
| KD | MAP2K2 | -95.87 | -95.83 | -98.49 | 39.01 | -43.99 | 27.13 | -95.9 | -99.22 | -98.77 | -60.87 | -99.21 |
| CP | vinorelbine | -95.9 | -61.1 | -98.95 | -95.61 | -12.87 | 38.39 | -96.52 | -96.2 | -99.65 | 46.62 | -98.48 |
| CP | diprotin-a | -95.91 | -95.91 | -70.2 | NA | NA | -18.89 | -98.44 | NA | NA | NA | -97.13 |
| KD | CAST | -95.95 | -88.06 | 97.76 | 53.73 | -99.26 | -98.35 | NA | 81.07 | -95.95 | -98.88 | -98.68 |
| CP | NSC-632839 | -95.98 | -75.25 | -98.61 | -98.5 | -82.67 | -62.01 | -97.41 | -65.38 | -99.84 | -96.75 | -95.21 |
| KD | KCNN4 | -95.98 | -45.13 | 43.32 | -99.34 | -99.95 | -93.39 | NA | -75.53 | -99.79 | -95.98 | -99.74 |

| | | | | | | | | | | | | |
|---|---|---|---|---|---|---|---|---|---|---|---|---|
| KD | GPSM2 | -96.03 | -98.13 | -47.92 | -85.19 | -94.47 | -99.41 | NA | NA | -98.58 | -36.55 | -97.59 |
| KD | ETV5 | -96.05 | 91.51 | -55.6 | 38.14 | -99.47 | -99.43 | NA | -97.31 | -96.05 | -38.92 | -99.79 |
| KD | GAS7 | -96.1 | -90.92 | -99.89 | -58.44 | NA | -94.49 | NA | -97.71 | -64.22 | -99.92 | -99.58 |
| KD | CDC7 | -96.14 | -37.9 | -26.5 | -99.98 | NA | NA | NA | -96.14 | -84.26 | -98.32 | -98.63 |
| CP | VU-0365114-2 | -96.15 | -99.78 | -99.43 | -36.81 | -96.15 | NA | -51.02 | -98.04 | 57.02 | -85.45 | -98.45 |
| CP | dihydroergotamine | -96.28 | -50.48 | -98.21 | 61.04 | -89.8 | -96.89 | -64.85 | -95.66 | -99.98 | -98.07 | -99.93 |
| KD | CARD11 | -96.52 | -96.34 | 91.89 | -98.78 | -90.81 | -98.87 | -52.66 | -31.15 | -96.71 | -99.29 | -99.75 |
| KD | MCM7 | -96.6 | -99.62 | -98.36 | 15.61 | -99.86 | -52.53 | -23.64 | -85.73 | -94.84 | -99.52 | -99.92 |
| CP | ruxolitinib | -96.66 | -75.45 | 20.69 | 14.47 | -98.26 | -99.89 | 22.27 | -99.23 | -95.05 | -99.88 | -99.86 |
| KD | FGFR4 | -96.78 | -97.59 | 99.42 | -93.68 | -99.84 | NA | NA | -84.81 | -95.97 | -99.96 | -99.4 |
| KD | GNAQ | -96.92 | -35.12 | -94.13 | -99.96 | -95.24 | -98.6 | NA | NA | -81.64 | -98.94 | -99.61 |
| KD | KSR2 | -96.95 | -96.95 | -35.95 | -29.2 | 60.33 | -97.08 | NA | 37.55 | -97.65 | -99.69 | -97.55 |
| KD | METRN | -97.09 | -99.83 | 24.46 | -98.52 | -95.66 | 51.55 | -84.48 | -99.96 | -99.96 | -92.76 | -99.95 |
| KD | APOC3 | -97.23 | -99.46 | -99.93 | -97.23 | NA | -63.62 | -99.07 | 28.87 | -39.49 | -20.55 | -99.61 |
| KD | HES5 | -97.25 | -69.37 | -28.69 | -99.4 | -60.83 | -37.7 | -98.22 | -99.96 | -97.25 | NA | -99.34 |
| KD | HOXA9 | -97.4 | -98.01 | NA | -96.79 | -92.59 | -82.5 | NA | -99.98 | -99.84 | -83.83 | -99.82 |
| KD | IGSF8 | -97.54 | -66.41 | -100 | -98.16 | -95.95 | -73.67 | -34.07 | -96.91 | -98.63 | -99.97 | -99.84 |
| CP | triciribine | -97.56 | -95.7 | -99.63 | -93.34 | 98.93 | -35.27 | -99.85 | -26.97 | -99.61 | -99.41 | -99.72 |
| KD | KBTBD2 | -97.7 | NA | -96.45 | -98.15 | -97.7 | -93.27 | NA | NA | 55.87 | -99.46 | -99.47 |
| KD | CAPN1 | -97.78 | 41.69 | 24.1 | -98.38 | -97.69 | -97.86 | -48.85 | -99.98 | -93.43 | -99.93 | -99.79 |
| KD | ARHGEF2 | -97.83 | 45.44 | -76.34 | -99.93 | -99.95 | -94.81 | NA | -89.65 | -97.83 | -99.47 | -99.87 |
| CP | emetine | -97.91 | -99.09 | -99.69 | -54.38 | -98.85 | -92.84 | -97.3 | -97.79 | -99.58 | -96.04 | -98.03 |
| CP | triciribine | -97.97 | -97.92 | 73 | -99.25 | 49.35 | -98.03 | -99.84 | 27.64 | -90.44 | -98.67 | -99.44 |
| KD | PCK2 | -98.06 | -38.64 | -98.15 | -98.06 | -69.62 | -47.87 | -99.97 | -99.94 | -97.53 | -98.05 | -99.87 |
| CP | MST-312 | -98.24 | -86.19 | -98.18 | -99.42 | -99.83 | -89 | -43.09 | -26.14 | -99.92 | -98.31 | -99.33 |
| KD | STUB1 | -98.3 | -98.3 | -20.37 | -98.17 | -99.22 | -98.46 | 55.94 | NA | -98.21 | -99.89 | -99.74 |
| KD | PCGF1 | -98.42 | -98.42 | -87.04 | 71.82 | -55.16 | -99.64 | NA | -99.83 | -99.31 | 37.15 | -99.92 |
| KD | BCL2L1 | -98.59 | -99.6 | -30.77 | -99.86 | -83.79 | 29.82 | NA | -98.59 | -32.19 | -98.71 | -99.92 |
| KD | RXRB | -98.6 | -90.88 | -100 | -99.79 | -98.6 | -99.95 | NA | -66.17 | 28.11 | -76.19 | -99.95 |

| | | | | | | | | | | | | |
|---|---|---|---|---|---|---|---|---|---|---|---|---|
| KD | ARF6 | -98.64 | -99.95 | 26.07 | -97.87 | -93.08 | -21.62 | 85.81 | -99.98 | -99.73 | -99.4 | -99.95 |
| OE | PPIA | -98.65 | 51.32 | -93.13 | -99.42 | 65.73 | -98.29 | -99 | -99.95 | 82.49 | -99.96 | -99.91 |
| KD | GAS6 | -98.81 | -99.13 | 46.01 | -98.86 | -99.93 | -98.77 | 59.08 | 38.71 | -99.89 | -71.92 | -99.79 |
| KD | LSP1 | -98.83 | -99.6 | NA | -99.88 | -98.83 | -99.93 | 56.89 | -71.12 | -97.01 | 42.38 | -99.92 |
| CP | GW-843682X | -98.98 | -99.84 | -95.66 | -99.76 | -99.58 | -97.72 | -97.86 | -75.84 | -99.95 | -98.38 | -99.61 |
| CP | kinetin-riboside | -99.23 | -99.45 | -77.29 | -92.7 | -99.48 | -99.26 | -99.87 | -99.88 | -97.11 | -98.45 | -99.19 |
| KD | PTK2B | -99.34 | -54.87 | -99.15 | -99.53 | NA | NA | -35.39 | -51.18 | -99.82 | -99.81 | -99.67 |
| KD | RTCD1 | -99.69 | -99.57 | -99.89 | -99.8 | -96.69 | -92.99 | -99.91 | 49.8 | -99.88 | -36.08 | -99.89 |
| KD | SMARCA2 | -99.85 | -26.1 | -100 | NA | -91.96 | -99.87 | NA | 63.61 | -99.98 | -99.82 | -99.95 |

CP, compound; KD, gene knock-down; OE, gene overexpression; PCL, perturbational class
